# A novel equilibrative nucleoside transporter 1 inhibitor alleviates Tau-mediated neurodegeneration

**DOI:** 10.1101/2020.10.25.334201

**Authors:** Ching-Pang Chang, Ya-Gin Chang, Pei-Yun Chuang, Thi Ngoc Anh Nguyen, Fang-Yi Chou, Sin-Jhong Cheng, Hui-Mei Chen, Lee-Way Jin, Kevin Carvalho, Vincent Huin, Luc Buée, David Blum, Yung-Feng Liao, Chun-Jung Lin, Yijuang Chern

**Affiliations:** Institute of Biomedical Sciences, Academia Sinica, Taipei, Taiwan; School of Pharmacy, National Taiwan University, Taipei, Taiwan; Neuroscience Program of Academia Sinica, Academia Sinica, Taipei, Taiwan; Department of Pathology and Laboratory Medicine, University of California Davis, Sacramento, CA, USA; Univ. Lille, Inserm, CHU Lille, U1172 - LilNCog - Lille Neuroscience & Cognition, F-59000 Lille, France; Alzheimer & Tauopathies, LabEx DISTALZ, LiCEND, F-59000 Lille, France; Institute of Cellular and Organismic Biology, Academia Sinica, Taipei, Taiwan

**Keywords:** Alzheimer’s disease, tauopathy, adenosine, AMPK, ENT1

## Abstract

Tau hyperphosphorylation favors the formation of neurofibrillary tangles and triggers the gradual loss of neuronal functions in tauopathies, including Alzheimer’s disease. Herein, we demonstrated that chronic treatment with an inhibitor (J4) of equilibrative nucleoside transporter 1 (ENT1), which plays a critical role in controlling adenosine homeostasis and purine metabolism in the brain, exerted beneficial effects in a mouse model of tauopathy (Thy-Tau22, Tau22). Chronic treatment with J4 improved spatial memory deficits, mitochondrial dysfunction, synaptic plasticity impairment, and gliosis. Immunofluorescence assays showed that J4 not only reduced Tau hyperphosphorylation but also normalized the reduction in mitochondrial mass and suppressed the abnormal activation of AMP-activated protein kinase (AMPK), a pathogenic feature that is also observed in the brains of patients with tauopathies. Given that AMPK is an important energy sensor, our findings suggest that energy dysfunction is associated with tauopathy and that J4 may exert its protective effect by improving energy homeostasis. Bulk RNA-seq analysis revealed that J4 also mitigated immune signature associated with Tau pathology including C1q upregulation and A1 astrocyte markers. Collectively, our findings suggest that identifying strategies for normalizing energy and neuroimmune dysfunctions in tauopathies through adenosinergic signaling modulation may pave the way for the development of treatments for Alzheimer’s disease.

## Introduction

Alzheimer’s disease is the most prominent neurodegenerative disease in aging societies, but there is no effective treatment (Cummings *et al.*, 2019). The major pathogenic hallmarks of Alzheimer’s disease include extracellular amyloid plaques (amyloid-beta, Aβ) and intracellular accumulation of neurofibrillary tangles made of hyperphosphorylated Tau. The latter is particularly known to be associated with neuritic dystrophy, synapse loss, and neuroinflammation, which lead to cognitive impairments (Ittner and Gotz, 2011; Brier *et al.*, 2016; Laurent *et al.*, 2018; Colin *et al.*, 2020). Phosphorylation of Tau can be modulated by more than 30 kinases and protein phosphatases (Sergeant *et al.*, 2008; Qian *et al.*, 2010), thus making it very sensitive to the environment including metabolic changes (Merkwirth *et al.*, 2012; Lee *et al.*, 2013).

Adenosine is an important homeostatic building block of many important metabolic pathways. It also serves as a neuromodulator that controls multiple functions (including neuroinflammation, blood-brain barrier permeability, neuronal transmission, and energy balance) through receptor-dependent and/or independent mechanisms in the central nervous system (Carman *et al.*, 2011; Chen *et al.*, 2014). The main sources of adenosine include ATP catabolism and the transmethylation pathway. In addition, the extracellular and intracellular adenosine levels are modulated through equilibrative nucleoside transporters (ENTs) and concentrative nucleoside transporters (Ham and Evans, 2012; Lee and Chern, 2014). Disruption of adenosine homeostasis in the brain has been implicated in sleep disorders, impaired cognition impairment and neurodegenerative diseases (Porkka-Heiskanen and Kalinchuk, 2011; Cunha, 2016). A reduction in adenosine levels has been observed in the frontal-, temporal-, and parietal-cortices of human Alzheimer’s disease brains (Alonso-Andres *et al.*, 2018). Abnormal expression of A_1_ and A_2A_ adenosine receptors has also been reported in the brain of patients (Albasanz *et al.*, 2008; Temido-Ferreira *et al.*, 2018) and mouse models with Alzheimer’s disease (Viana da Silva *et al.*, 2016; Faivre *et al.*, 2018; Lee *et al.*, 2018; Silva *et al.*, 2018), suggesting that the adenosinergic system is dysregulated in Alzheimer’s disease. However, the link between adenosine homeostasis and Tau pathology remains ill-defined.

AMP-activated protein kinase (AMPK) is another homeostatic energy sensor that controls the balance between anabolic and catabolic processes in cells (Herzig and Shaw, 2018). In the presence of stress (e.g., an elevated AMP/ATP ratio, high reactive oxygen species (ROS) levels, or mitochondrial dysfunction), AMPK can be activated by being phosphorylated at Thr^172^ on the a subunit of AMPK (Turnley *et al.*, 1999). Under adverse conditions when the extracellular adenosine concentration is elevated, transport of adenosine into cells enhances the cellular level of AMP, alters the AMP/ATP ratio, and subsequently activates AMPK (Pastor-Anglada *et al.*, 2007; Medina-Pulido *et al.*, 2013). Previous studies have associated energy deprivation and AMPK activation with abnormal Tau phosphorylation in the brains of patients and mice with Alzheimer’s disease (Vingtdeux *et al.*, 2011; Ma *et al.*, 2014; Lauretti *et al.*, 2017; Fang *et al.*, 2019).

We previously demonstrated the protective effects of adenosine augmentation against Alzheimer’s disease-like amyloid pathology by treating an Alzheimer’s disease mouse model (APPswe/PS1dE9) with an orally active and BBB-permeable blocker (J4) of ENT1. J4 is a competitive inhibitor of ENT1 with a Ki value of 0.05 μM (Lee *et al.*, 2018) and promotes the elevation of extracellular adenosine tone. In the present study, we investigated whether J4 exerts a protective effect against Tau pathology and associated cognitive deficits using the Thy-Tau22 (Tau22) model, which progressively develops hippocampal Tau pathology and memory deficits (Schindowski *et al.*, 2006). We demonstrated that chronic treatment with J4 mitigates the development of hippocampal Tau pathologies (including synapse loss, mitochondrial dysfunction, and neuroinflammation). These beneficial effects of J4 are particularly ascribed to its ability to suppress AMPK overactivation in the hippocampi of Tau22 mice, suggesting that it normalizes energy dysfunction caused by pathogenic Tau.

## Materials and Methods

### Animals and drug administration

Thy-Tau22 (Tau22) mice ((B6.Cg-Tg(Thy1-MAPT)22Schd) expressing mutant human 4R Tau (G272V and P301S) driven by a neuron-specific promoter (Thy1.2) and were maintained on the C57BL/6J background (Schindowski *et al.*, 2006). All animal studies were conducted following the protocols approved by the Institutional Animal Care and Utilization Committee (IACUC, Academia Sinica, Taiwan). All mice were housed in ventilated cages (IVC) with freely accessible water and chow diet (LabDiet®, San Antonio, TX, USA), and kept under controlled lighting condition with 12:12 hrs light/dark cycle at the Institute of Biomedical Sciences Animal Care Facility (Taipei, Taiwan). To investigate the effect of J4, male Tau22 mice and their littermate controls were randomly allocated to experimental groups that treated with the J4 (0.02 mg/ml in 1% HPβCD; designated TauJ and WTJ mice) or vehicle (1% HPβCD; designated TauC and WTC mice) continuously in drinking water for 7 months from the age of 3 to 10 months. Males were chosen for the following experiments. No sexual dimorphism was reported in the Tau22 strain (Laurent *et al.*, 2016).

### Human cases

For immunofluorescence analysis, a total of 18 post-mortem Human posterior hippocampal specimens: six normal subjects, six Alzheimer’s disease, and six FTD-Tau (CBD, PSP, and Pick’s), were obtained from the UC Davis Alzheimer’s Disease Center (USA).

For mRNA analysis, a total of 55 post-mortem Human brain samples (Brodmann area 10 prefrontal cortex or temporal cortex): 13 normal subjects, 19 Alzheimer’s Disease, and 23 FTD-Tau (CBD, P301L, PSP, and Pick’s), were obtained from the brain banks of Lille, Paris, and Geneva. Participants and methods have been described previously (Huin *et al.*, 2016; Carvalho *et al.*, 2019). Fresh frozen grey matter tissue (about 100 mg) retrieved at autopsy and stored at −80°C was used for mRNA analysis. All the brain samples used for RT-qPCR analyses had an RNA integrity number ≥ 5. Detailed information on the normal subjects, Alzheimer’s Disease patients, and FTD-Tau patients from which the specimens used in the present study were obtained is listed in Supplementary Table 1.

### Morris water maze

Spatial memory and cognitive flexibility of mice aged 10-11 months were evaluated using the Morris water maze test as described with slight modifications (Chang *et al.*, 2016). A detailed procedure is given in the supplementary material.

### Electrophysiological study

Mice aged 10-11 months were used for electrophysiology approaches. All electrophysiology studies were performed at the electrophysiology core facility (Neuroscience Program of Academia Sinica, Taipei, Taiwan). After rapid decapitation, the hippocampus was quickly dissected out and immersed in the ice-cold artificial cerebrospinal fluid (ACSF; 119 mM NaCl, 2.5 mM KCl, 2.5 mM CaCl_2_, 1.3 mM MgSO_4_, 1 mM NaH_2_PO_4_, 26.2 mM NaHCO_3_, and 11 mM glucose) oxygenated with 95% O_2_ and 5% CO_2_. Transverse hippocampal slices of 450 μm thickness were prepared with a DSK Microslicer (DTK-1000, Osaka, Japan), recovered in oxygenated ACSF at RT for 3 hrs, and then recorded the field excitatory postsynaptic potentials (fEPSPs). Input-output curves, paired-pulse facilitation (PPF), and long-term depression (LTD) at the Schaffer collateral-CA1 synapses were evaluated as described elsewhere (Van der Jeugd *et al.*, 2011). A detailed procedure is given in the supplementary material.

### RNA extraction, cDNA synthesis, and quantitative PCR

RNA isolation, complementary DNA (cDNA) synthesis, quantitative PCR (qPCR) of mouse and human brain tissue were performed as described previously (Lee *et al.*, 2018; Carvalho *et al.*, 2019). The reference genes for mouse and human brain tissue are GAPDH and β-Actin (ACTB), respectively. The PCR primers and probes used in this study are shown in Supplementary Tables 2 and 3. A detailed procedure is given in the supplementary material.

### RNA sequencing (RNA-seq)

Total RNA samples (3 μg per sample) extracted from the hippocampus with RIN values greater than 8 were subjected to RNA-seq analysis. The RNA library construction, sequencing (150-bp paired-end reads), alignments (mm10 version of the mouse genome, Ensembl release 93), and gene expression analysis (fragments per kilobase per million, FPKM) were carried out by Welgene Biotech (Taipei, Taiwan). A detailed procedure is given in the supplementary material. To identify the differentially expressed (DE) genes in different groups, cutoff criteria (absolute log2 fold change ≥ 0.32, *p*<0.05) were used. A volcano plot and heatmap of the DE genes were drawn by using Instant Clue (Nolte *et al.*, 2018) and Morpheus software (https://software.broadinstitute.org/morpheus/), respectively. The Gene Ontology (GO) and the Kyoto Encyclopedia of Genes and Genomes (KEGG) pathways of DE genes were analyzed by using the Database for Annotation, Visualization and Integrated Discovery (DAVID 6.7) (Huang da *et al.*, 2009a, b). Ingenuity Pathway Analysis (IPA) software (Qiagen, CA, USA) was then used for the identification of signaling pathway(s) or upstream regulator(s) related to the DE genes.

### Immunoblotting

Total proteins (20 μg) were used for western blot analyses, the immunosignals were detected using enhanced chemiluminescence (ECL) reagent (PerkinElmer, MA, USA). The primary antibodies used in this study are listed in Supplementary Table 4. Detailed biochemical procedures are provided in the Supplementary material.

### Immunohistochemical staining

For mouse brain tissue, coronal brain sections (20 μm) were prepared and used for immunostaining as previously described (Lee *et al.*, 2018). For human brain tissue, formalin-fixed, paraffin-embedded human brain slices (6 μm) were deparaffinized, rehydrated, and immersed in 1X citrate buffer (C9999, Sigma-Aldrich, St. Louis, MO, USA) to expose the antigenic sites. The human brain slices were subjected to immunofluorescence staining and the autofluorescence signal was blocked by 0.1% (w/v) Sudan Black B (199664, Sigma) in 70% ethanol. The images were captured by the LSM 780 confocal microscope (Carl Zeiss, Germany) and then analyzed with MetaMorph software (Universal Imaging, PA, USA). The primary antibodies used in this study are listed in Supplementary Table 4. Detailed immunostaining procedures are provided in the Supplementary material.

### Statistics

The experimental condition was blinded to investigators during behavioral and electrophysiological experiments. The data are expressed as the mean ± S.E.M. All statistical analyses were performed using GraphPad Prism Software (La Jolla, CA, USA). Two-tailed unpaired Student’s *t*-test was used to compare the difference between the two groups. One-way or two-way ANOVA followed by Tukey’s multiple comparisons test was used for comparison of multiple groups. Differences were considered statistically significant when *p* < 0.05.

### Data availability

The data that support the findings of this study are available on request to the corresponding author.

## Results

### Chronic J4 treatment prevents impairment of spatial learning and memory of Tau22 mice

To evaluate the effect of J4 on Tau22 mice, we treated these mice with J4 (0.02 mg/ml in 1% HPβCD) or vehicle (1% HPβCD) in drinking water for 7 months beginning at 3 months of age. Based on the average drinking water intake of each mouse (4.9 ± 0.4 ml/day; n = 20), the average daily intake level of J4 was 3.06 ± 0.28 mg/kg. The inclusion of J4 in drinking water did not affect the amount of water intake (data not shown). No adverse events (e.g., weight loss, vomiting, or sudden death) were observed during experiments. Behavioral and biochemical analyses of these mice were conducted when they were 10-11 months old.

The spatial memory and cognitive flexibility of Tau22 mice at the age of 10 months were evaluated using the Morris water maze (MWM) task. In the acquisition-learning phase (Day 1-Day 5), vehicle-treated Tau22 mice (TauC) spent a longer time locating the hidden platform than vehicle-treated WT mice (WTC) (Fig. 1a, \**p* < 0.05, versus WTC mice, two-way ANOVA), showing that Tau22 mice exhibited spatial learning impairment, as previously reported (Schindowski *et al.*, 2006). J4 treatment significantly prevented spatial learning deficits in Tau22 mice (Fig. 1a, ^#^*p* < 0.05, versus TauC mice, two-way ANOVA) without affecting wild-type mice. We next performed the probe test on Day 8 to assess spatial memory. Vehicle- and J4-treated WT mice showed a preference for searching the hidden platform in the target quadrant (T) over the nontarget quadrants (NTs) (Fig. 1b, \**p* < 0.05, versus NTs, two-tailed Student’s *t*-test). As expected, TauC mice did not prefer the T quadrant over the NT quadrants (Fig. 1b), indicating that the spatial memory of Tau22 mice was impaired. The impaired memory was normalized by chronic J4 treatment (Fig. 1b, \**p* < 0.05, versus NTs, twotailed Student’s *t*-test).

**Fig. 1.**
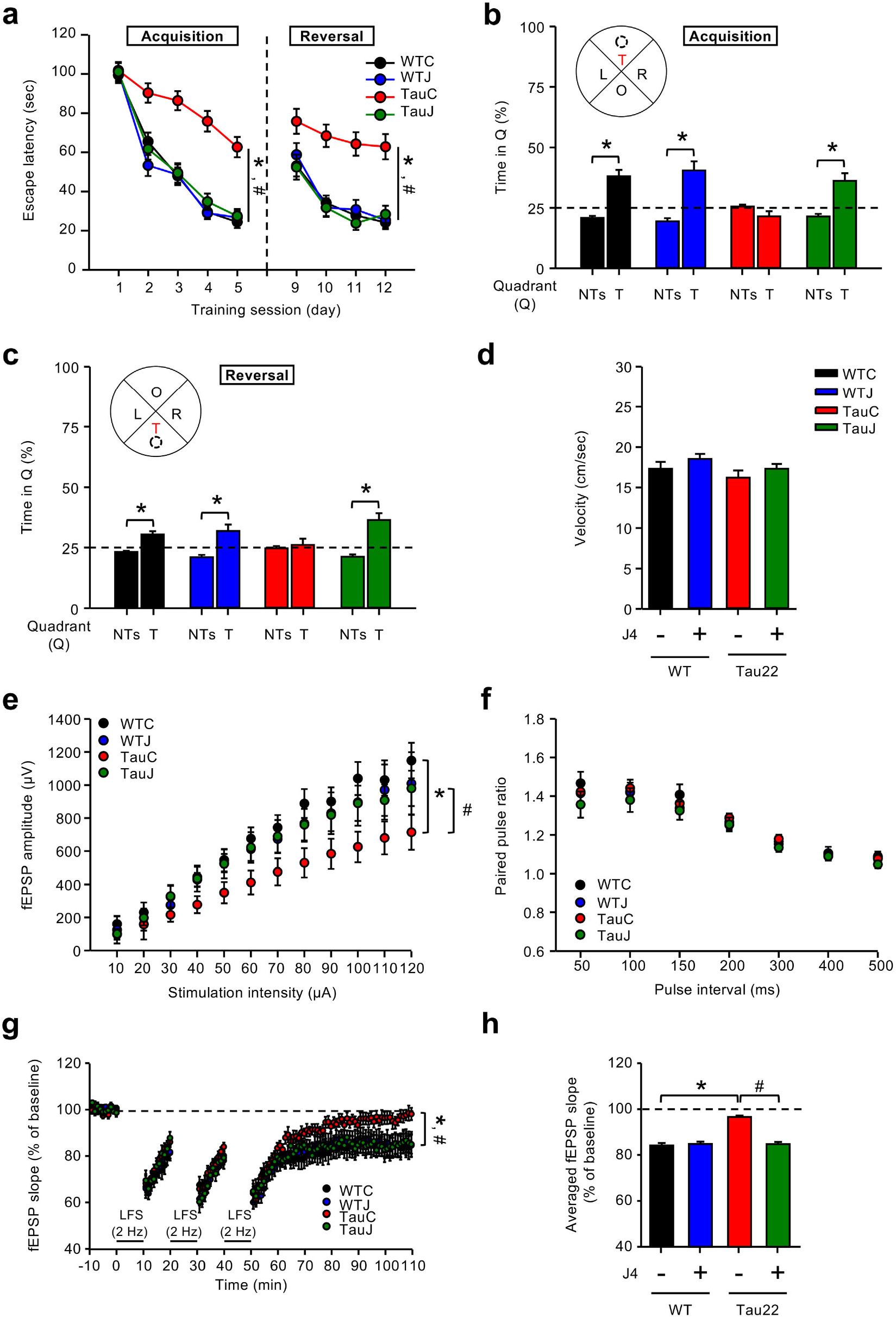
Chronic treatment with J4 alleviates the impairment of spatial memory and hippocampal CA1 LTD in Tau22 mice. **(a-d)** Mice were treated as indicated (control WT mice, WTC, black; J4-treated WT mice, WTJ, blue; control Tau22 mice, TauC, red; and J4-treated Tau22 mice, TauJ, green; n= 12-20 in each group) from the age of 3-10 months. The Morris water maze (MWM) was used to assess spatial learning and memory. **a.** In the acquisition-learning phase, learning trials with a hidden platform were performed for five days (Day 1-Day 5); in the reversal-learning phase, learning trials with a hidden platform (opposite quadrant) were performed for an additional four days (Day 9-Day 12). *\*p* < 0.05 versus the WTC group; ^#^*p* < 0.05 versus the TauC group; two-way ANOVA. **b-c.** For the spatial memory and spatial reversal memory, probe tests were performed 72 hrs after the last trial of **(b)** the acquisition-learning phase (Day 8) and **(c)** the reversal learning phase (Day 15). Performance in the probe test is presented as the averaged percentage of time spent in the T (target quadrant) and NTs (nontarget quadrants). The horizontal dashed lines shown in **(b)** and **(c)** represents the expected percentage of time spent in each quadrant by chance (25%) in the probe test. \**p* < 0.05 versus the nontarget quadrants; two-tailed Student’s *t*-test. **d.** The swimming speed (cm/s) of the animals in the probe test was measured, and no significant difference was identified (one-way ANOVA). The data are expressed as the means ± SEM. **(e-h)** Hippocampal slices were prepared from mice subjected to different treatment groups (WTC, black; WTJ, blue; TauC, red; and TauJ, green) from the age of 3-10 months. **e.** The input/output relationship curve at the Schaffer collateral-CA1 synapses. The mean fEPSP responses to stimuli of increasing strengths (from 10 μA to 120 μA) were recorded from mice subjected to different treatments (WTC, n=7 slices from 6 mice; WTJ, n=3 slices from 3 mice; TauC, n=9 slices from 5 mice; TauJ, n=7 slices from 5 mice). **f.** Paired-pulse facilitation at the Schaffer collateral-CA1 synapses. Averaged paired-pulse ratios in response to stimuli at different interstimulus intervals (from 50 ms to 500 ms) were recorded from mice subjected to different treatments (WTC, n=6 slices from 3 mice; WTJ, n=11 slices from 3 mice; TauC, n=13 slices from 4 mice; TauJ, n=11 slices from 3 mice). **(g-h).** Long-term depression (LTD) at the Schaffer collateral-CA1 synapses. LTD was induced by 3 trains of 1200 pulses at 2 Hz LFS in the CA1 region. **(g)** The average fEPSP slopes of mice subjected to different treatments (WTC, n=18 slices from 7 mice; WTJ, n=15 slices from 5 mice; TauC, n=14 slices from 8 mice; TauJ, n=12 slices from 6 mice) were calculated. \**p* < 0.05 versus the WTC group; ^#^*p* < 0.05 versus the TauC group, two-way ANOVA. **(h)** Quantification results of the mean fEPSP slopes during the last 10 min of the steady-state period of the recording after LTD induction. The data are expressed as means ± S.E.M. \**p* < 0.05 versus the WTC group; ^#^*p* < 0.05 versus the TauC group, one-way ANOVA.

To assess cognitive flexibility, we further examined the spatial reversal learning of WT and Tau22 mice treated with or without J4 for an additional four consecutive days. In the reversal-learning phase (Day 9-Day 12), the hidden platform was relocated to the opposite quadrant and the escape latency was recorded. The ability of TauC mice to find the new location of the hidden platform was impaired compared with that of WTC mice (Fig. 1a, *\*p* < 0.05, versus WTC mice, two-way ANOVA). The impaired spatial reversal learning of Tau22 mice was also improved by J4 (Fig. 1a, ^#^*p* < 0.05, versus TauC mice, two-way ANOVA). To evaluate spatial reversal memory, each mouse was subjected to the probe test 72 hrs after the last trial of reversal learning trial on Day 15. WTC and WTJ mice both spent more time in the new target quadrant (T) than in the nontarget quadrants (NTs) (Fig. 1c, \**p* < 0.05, versus NTs, two-tailed Student’s *t*-test). Conversely, the TauC mice failed to distinguish the T and NTs (Fig. 1c, *p* = 0.62, versus NTs, two-tailed Student’s *t*-test). Most importantly, chronic J4 treatment prevented the impairment of cognitive flexibility in Tau22 mice (Fig. 1c, \**p* < 0.05, versus NTs, twotailed Student’s *t*-test). No difference in the swimming speed was observed among the groups tested (Fig. 1d). Together, our data indicate that chronic treatment with J4 significantly improves the spatial memory and spatial reversal memory of Tau22 mice.

### Chronic J4 treatment normalizes the impairment of hippocampal CA1 LTD in Tau22 mice

We next investigated whether chronic J4 treatment affects the synaptic plasticity in Tau22 mice at the age of 10-11 months. The basal transmission of the hippocampal CA3-CA1 network was first determined based on the input-output relationship. The vehicle-treated Tau22 mice showed decreased basal synaptic transmission compared to that of the vehicle-treated WT mice (Fig. 1e, \**p* < 0.05, versus WTC mice, two-way ANOVA). Under the strongest stimulation (120 μA) used, the mean fEPSP amplitude (μV) of TauC mice was 38% lower than that of WTC mice. J4 alleviated the abnormal synaptic transmission in Tau22 mice (Fig. 1e, ^#^*p* < 0.05, versus TauC mice, two-way ANOVA), but did not impact than in WT mice. Presynaptic neurotransmitter release in the hippocampus, as determined by the paired-pulse facilitation (PPF) response assay, was comparable between genotypes (Tau22 versus WT mice) and treatment groups (J4 versus vehicle) (Fig. 1f, *p* > 0.05, two-way ANOVA), suggesting that presynaptic plasticity was unaffected in Tau22 mice.

Previous studies have demonstrated that Tau22 mice exhibit impaired long-term depression (LTD) but normal long-term potentiation (LTP) at the Schaffer collateral-CA1 synapses in the hippocampus (Schindowski *et al.*, 2006). As shown in Fig. 1g, LTD was maintained in WTC mice but not in TauC mice (\**p* < 0.05, versus WTC mice, two-way ANOVA). This impairment in LTD was prevented by chronic J4 treatment (^#^*p* < 0.05, versus TauC mice, two-way ANOVA). The average LTD magnitude during the last 10 min of recording was quantified and shown in Fig. 1h (\**p* < 0.05, versus WTC mice; ^#^*p* < 0.05, versus TauC mice, one-way ANOVA). Collectively, these results demonstrate that chronic treatment with J4 normalizes the impaired basal synaptic transmission and LTD at Schaffer collateral synapses in the hippocampi of Tau22 mice without affecting those in the hippocampi of WT mice.

### Chronic J4 treatment reduces Tau hyperphosphorylation in the hippocampi of Tau22 mice

Tau hyperphosphorylation and subsequent misfolding are pathophysiological triggers in Tau22 mice (Schindowski *et al.*, 2006). To examine the effect of J4 on Tau phosphorylation in the hippocampi of Tau22 mice, we first performed a differential peptide labeling (iTRAQ) of hippocampal proteins and analyzed the results using LC-MS/MS-based proteomics. Phosphopeptides covering 30 phosphorylation sites of human Tau were identified (Supplementary Fig. 1). Chronic treatment with J4 reduced phosphorylation at 15 phosphorylation sites. The J4-mediated reduction in phosphorylation of Tau at Alzheimer’s Disease-associated phosphorylation sites in the hippocampi of Tau22 mice was further assessed using Western blot and immunofluorescence analysis using antibodies raised against hyperphosphorylated (pSer181, pSer199, pSer202/Thr205 = AT8, pSer262, pSer396, and pSer422) and misfolded (pThr212/Ser214 = AT100; MC1) Tau. Chronic treatment with J4 significantly reduced the phosphorylation levels of Tau at the sites tested except for pSer181 (Figs. 2a-2d, ^#^*p* < 0.05, versus TauC mice, two-tailed Student’s *t*-test). No effect of J4 on the protein and transcript levels of human Tau was observed, demonstrating that J4 did not modulate the Thy-1 promoter directly (Supplementary Fig. 2, *p* > 0.05, versus TauC mice, two-tailed Student’s *t*-test). Importantly, the levels of misfolded Tau, which was detected with AT100 and MC1 (Jicha *et al.*, 1997; Mellone *et al.*, 2013), were also reduced by J4 (Figs. 2a, 2c and 2d, ^#^*p* < 0.05, versus TauC mice, two-tailed Student’s *t*-test). These findings collectively suggest that chronic treatment with J4 reduces the levels of hyperphosphorylated and misfolded Tau in the hippocampi of Tau22 mice.

**Fig. 2.**
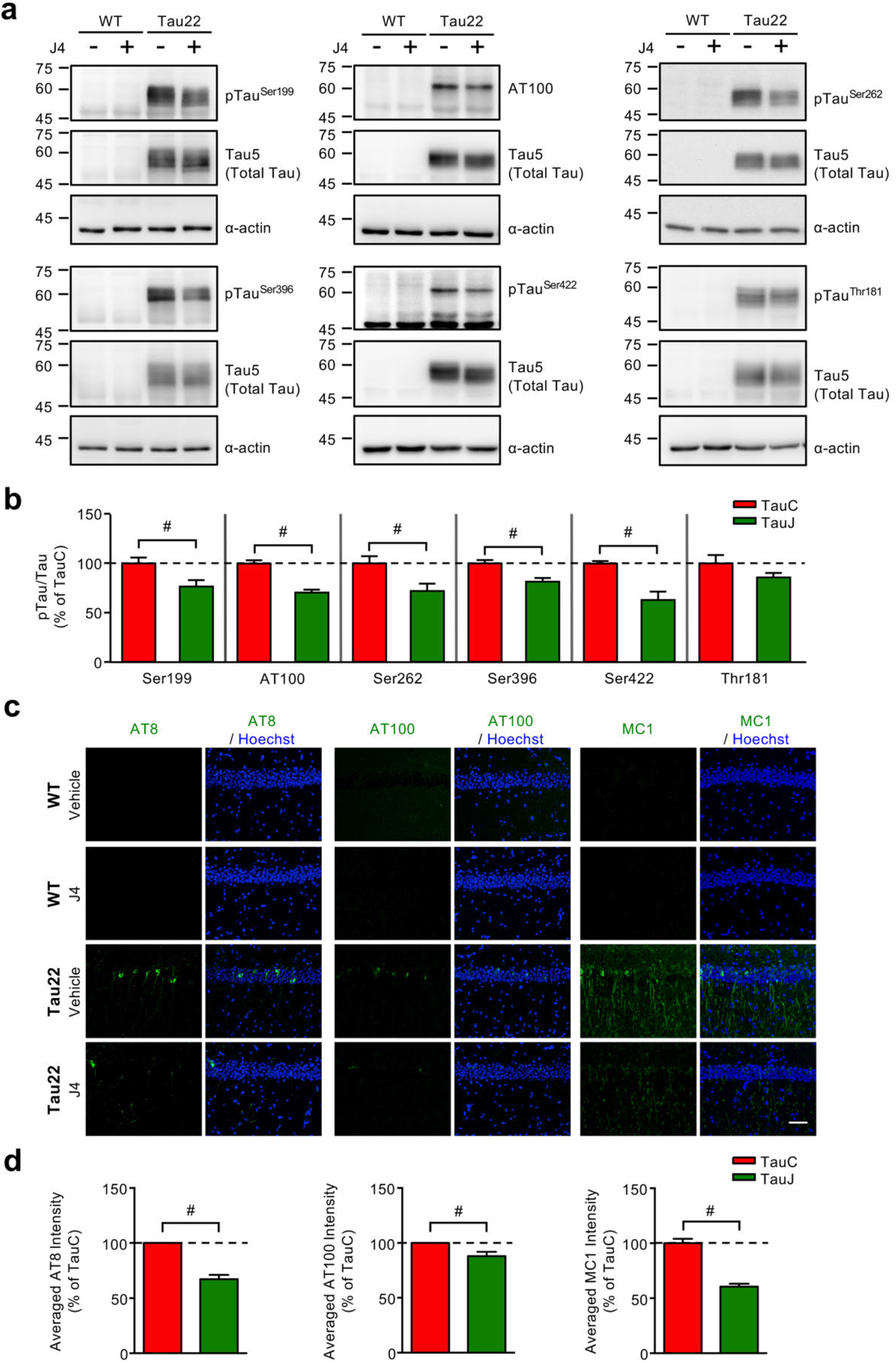
Chronic J4 treatment decreases hyperphosphorylated and misfolded human Tau levels in the hippocampi of Tau22 mice. Mice were treated as indicated (control WT mice, WTC, black; J4-treated WT mice, WTJ, blue; control Tau22 mice, TauC, red; and J4-treated Tau22 mice, TauJ, green; n = 3-4 in each group) from the age of 3-11 months. **(a-b)** Total hippocampal lysates (20 μg per lane) were harvested and subjected to WB analysis. **(a)** The protein expression levels of phospho-tau (pSer181, pSer199, pThr212/Ser214, pSer262, pSer396, and pSer422) and total tau were detected using the indicated antibodies. α-Actin was used as a loading control. Protein expression levels and phosphorylation levels were quantified and are shown in **(b)**. **(c-d)** Hippocampal sections (20 μm) were prepared from mice with different treatment groups (WTC, black; WTJ, blue; TauC, red; and TauJ, green; n = 3-5 in each group) from the age of 3-11 months and subjected to IHC staining. **(c)** The levels of hyperphosphorylated tau and misfolded tau in the hippocampus were evaluated by staining with the indicated antibodies (AT8 for Ser202/Thr205, green; AT100 for Thr212/Ser214, green; MC1 for conformational changed tau, green), and the quantification results of the average hyperphosphorylated tau intensity and averaged MC1 intensity are shown in **(d)**. Scale bar, 50 μm. The data are expressed as the mean ± S.E.M. ^#^*p* < 0.05, versus the TauC group, two-tailed Student’s *t*-test.

### Chronic J4 treatment reduces the AMPK activation in the hippocampus of Tau22 mice

To assess whether specific pathway(s) that may contribute to Tau phosphorylation are altered by J4 treatment in the hippocampi of Tau22 mice, we next examined the phosphorylation level of 17 kinases and signaling molecules by using the MAPK Phosphorylation Array (RayBiotech, GA, USA). Only minor or no change in the phosphorylation/activation of the kinases tested was found between Tau22 mice and WT mice (Supplementary Fig. 3). We also examined the expression and activation of PP2A, the major tau phosphatase (Martin *et al.*, 2013), by determining the total protein (i.e., PP2A-C), the Tyr^307^ phosphorylation, and the demethylation levels. No marked alterations were found between WT and Tau22 mice. Additionally, J4 did not exert a significant effect on these levels (Supplementary Fig. 4).

Because adenosine homeostasis has been implicated in the regulation of AMPK activation (Pastor-Anglada *et al.*, 2007; Medina-Pulido *et al.*, 2013; Sayama *et al.*, 2020), and since AMPK directly phosphorylates Tau (Domise *et al.*, 2016), we next evaluated the activation/phosphorylation of AMPK (Thr^172^, designated pAMPK) and Tau phosphorylation in the hippocampi of postmortem Alzheimer’s Disease and FTLD-Tau patients as well as Tau22 animals. Immunofluorescence staining revealed that, in the posterior hippocampal sections from Alzheimer’s Disease and FTD-Tau patients, pAMPK was detected in neurons that contained phosphorylated Tau (Fig. 3a and Supplementary Fig 5). Similarly, elevated phosphorylation of AMPK was observed in neurons containing phosphorylated Tau in the CA1 region in Tau22 mice, but rarely in those in wild-type mice (Figs. 3b and 3c). Chronic J4 treatment decreased the levels of pAMPK and pTau in Tau22 mice compared to WT mice (Figs. 3b and 3c, \**p* < 0.05, versus WTC mice; ^#^*p* < 0.05, versus TauC mice, one-way ANOVA; Figs. 3b and 3d, ^#^*p* < 0.05, versus TauC, two-tailed Student’s *t*-test).

**Fig. 3.**
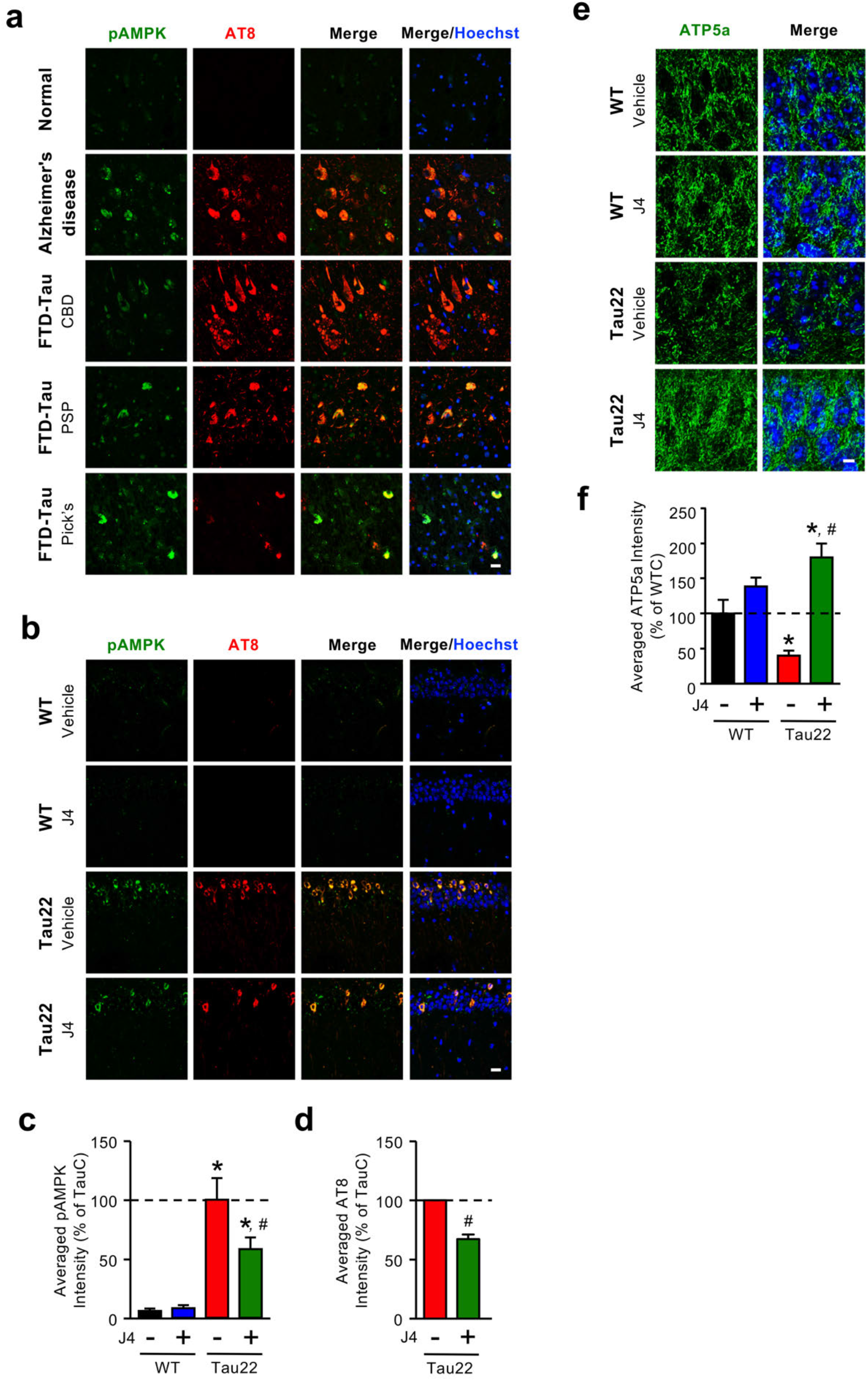
Chronic J4 treatment decreases AMPK activation and rescues mitochondrial abnormalities in the hippocampi of Tau22 mice. **(a)** Posterior hippocampal sections (6 μm) from normal subjects and Alzheimer’s Disease and FTD-Tau (CBD, PSP, and Pick’s disease) patients were subjected to IHC staining. The levels of phospho-AMPK and hyperphosphorylated tau were evaluated by staining with the indicated antibodies (pAMPK^Thr172^, green; AT8 for pTau^Ser202/Thr205^, red). **(b-f)** Mice were treated as indicated (control WT mice, WTC; J4-treated WT mice, WTJ; control Tau22 mice, TauC; and J4-treated Tau22 mice, TauJn; n= 5-7 in each group) from the age of 3-11 months. Hippocampal sections (20 μm) were prepared and subjected to IHC staining using the indicated antibodies (pAMPK^Thr172^, green; AT8 for pTau^Ser202/Thr205^, red), and the staining was quantified **(c, d)**. Scale bar, 20 μm. **(e-f)** The level of the mitochondrial marker ATP5a was evaluated by staining with an anti-ATP5a antibody (**e**, green) and quantified **(f;** WTC, black; WTJ, blue; TauC, red; TauJ, green, n = 3 in each group). Scale bar, 5 μm. The data are expressed as the mean ± S.E.M. \**p* < 0.05 versus the WTC group; ^#^*p* < 0.05, versus the TauC group, one-way ANOVA.

Since AMPK is a critical energy sensor and Tau has been implicated in mitochondrial dysfunction (DuBoff *et al.*, 2012), we next assessed mitochondrial mass by immunohistochemical staining using an antibody against ATP5A, a component of complex V (Cha *et al.*, 2015). Consistent with abnormal AMPK activation, TauC mice exhibited less ATP5A-positive mitochondrial mass in the hippocampus than WTC mice (Fig. 3e). J4 treatment ameliorated the loss of mitochondria in Tau22 mice (Fig. 3e and 3f, \**p* < 0.05, versus WTC; ^#^*p* < 0.05, versus TauC, one-way ANOVA) as it normalized the abnormal AMPK activation (Figs. 3b and 3c). These findings suggest that the blockade of ENT1 by J4 normalized Tau-associated energy dysfunction.

### Transcriptomic signature associated with the beneficial effect of J4 in the hippocampi of Tau22 mice

To gain mechanistic insight into how J4 mediates beneficial effects in Tau22 mice, we performed transcriptional profiling of the hippocampus using RNA-seq analysis. A total of 1441 differentially expressed (DE) genes (950 upregulated and 491 downregulated) between TauC and WTC mice were identified (Fig. 4a, TauC mice versus WTC mice, *p* < 0.05 and absolute log2 fold-change ≥ 0.32). Analyses of these DE genes by Ingenuity Pathway Analysis (IPA) revealed multiple canonical pathways (including neuroinflammation signaling pathway, dendritic cell maturation, TREM1 signaling) that were earlier found to be altered in several Alzheimer’s Disease mouse models (e.g. APP/PS1 and Tg4510; Supplementary Fig. 6; (Polito *et al.*, 2014; Benito *et al.*, 2015; Cummings *et al.*, 2015; Matarin *et al.*, 2015; Martinez Hernandez *et al.*, 2018; Sierksma *et al.*, 2020)) including Tau22 mice (Ising *et al.*, 2019). Interestingly, between the TauJ and WTC groups, a total of 1304 DE genes (417 upregulated and 887 downregulated, Fig. 4b; TauJ mice versus WTC mice, *p* < 0.05, absolute log2 fold-change ≥ 0.32) were also identified. Gene ontology (GO; Fig. 4c, FDR < 0.05, Benjamini *p* < 0.01) and Kyoto Encyclopedia of Genes and Genomes (KEGG; Fig. 4d, FDR < 0.05, Benjamini *p* < 0.01) analyses were conducted, and the DE genes between the TauC group vs WTC group were found to be associated with the immune response and transcriptional machinery. Interestingly, most of these pathways were not enriched in the DE genes between the TauJ group vs WTC group (Figs. 4c and 4d), suggesting that chronic J4 treatment normalized multiple pathways that were aberrantly regulated by pathogenic Tau.

**Fig. 4.**
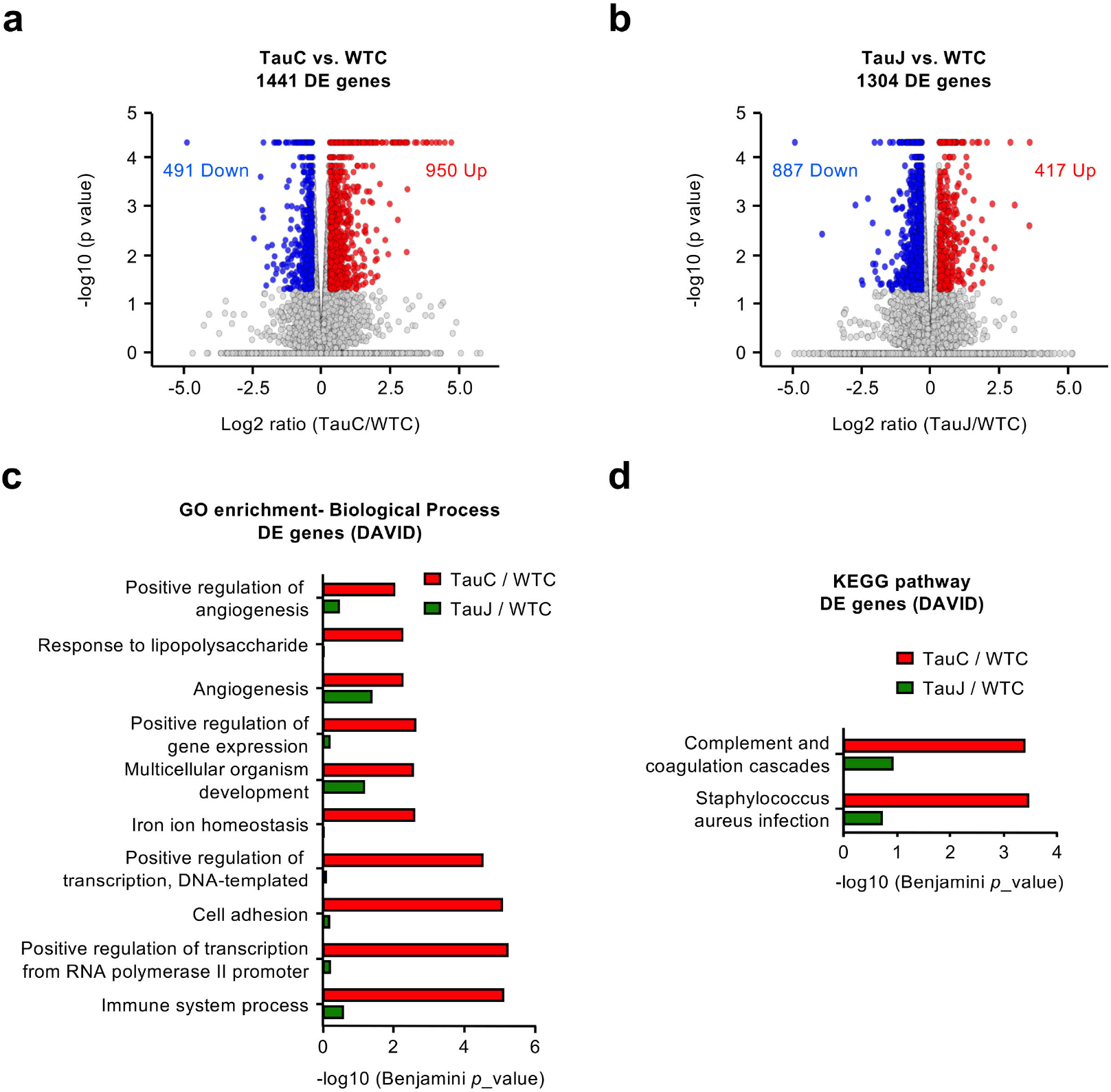
Chronic J4 treatment ameliorates the expression of tauopathy associated genes in the hippocampi of Tau22 mice. Mice were treated as indicated (control WT mice, WTC; control Tau22 mice, TauC; and J4-treated Tau22 mice, TauJ; n= 3 in each group) from the age of 3-10 months. The hippocampus was carefully removed for RNA-seq analysis. Volcano plot of DE genes in the **(a)** TauC group vs. WTC group and **(b)** TauJ group vs. WTC group. **(c)** GO enrichment analysis of DE genes between the TauC and WTC groups (red bar) and TauJ and WTC groups (green bar). **(d)** KEGG pathway analysis of DE genes between the TauC and WTC groups (red bar) and TauJ and WTC groups (green bar). In the volcano plot, the significantly upregulated and downregulated DE genes (absolute log2 ratio ≥ 0.32; *p* < 0.05) are shown in red and blue, respectively.

We also analyzed the effect of J4 on Tau22 mice. A total of 1239 DE genes were identified (289 upregulated and 950 downregulated; TauJ mice versus TauC mice, *p* < 0.05 and absolute log2 fold-change ≥ 0.32; Fig. 5a). J4 normalized the expression of 436 of the 950 upregulated DE genes (Fig. 5b) and 85 of the 491 downregulated DE genes (data not shown) between the TauC group and WTC group. The effect of J4 on the downregulated DE genes was not assessed further because no pathway was identified by GO and KEGG enrichment analysis (data not shown). For the upregulated DE genes with expression levels that were normalized by J4 (Fig. 5b), glycoprotein was identified by the UniProtKB keywords (UP_KEYWORDS, FDR < 0.05, Benjamini *p* < 0.01) (Fig. 5c). Further analyses revealed that these glycoprotein-related DE genes were enriched in pathways related to the immune response (Fig. 5d, GO analysis; Fig. 5e, KEGG analysis). Collectively, bulk RNAseq analysis suggested that chronic J4 treatment had a broad impact on Alzheimer’s Disease-related signaling molecules and pathways in the hippocampi of Tau22 mice, with a special focus on the inflammatory response that is evoked by pathogenic Tau and thought to contribute significantly to pathological development of tauopathies via a vicious circle.

**Fig. 5.**
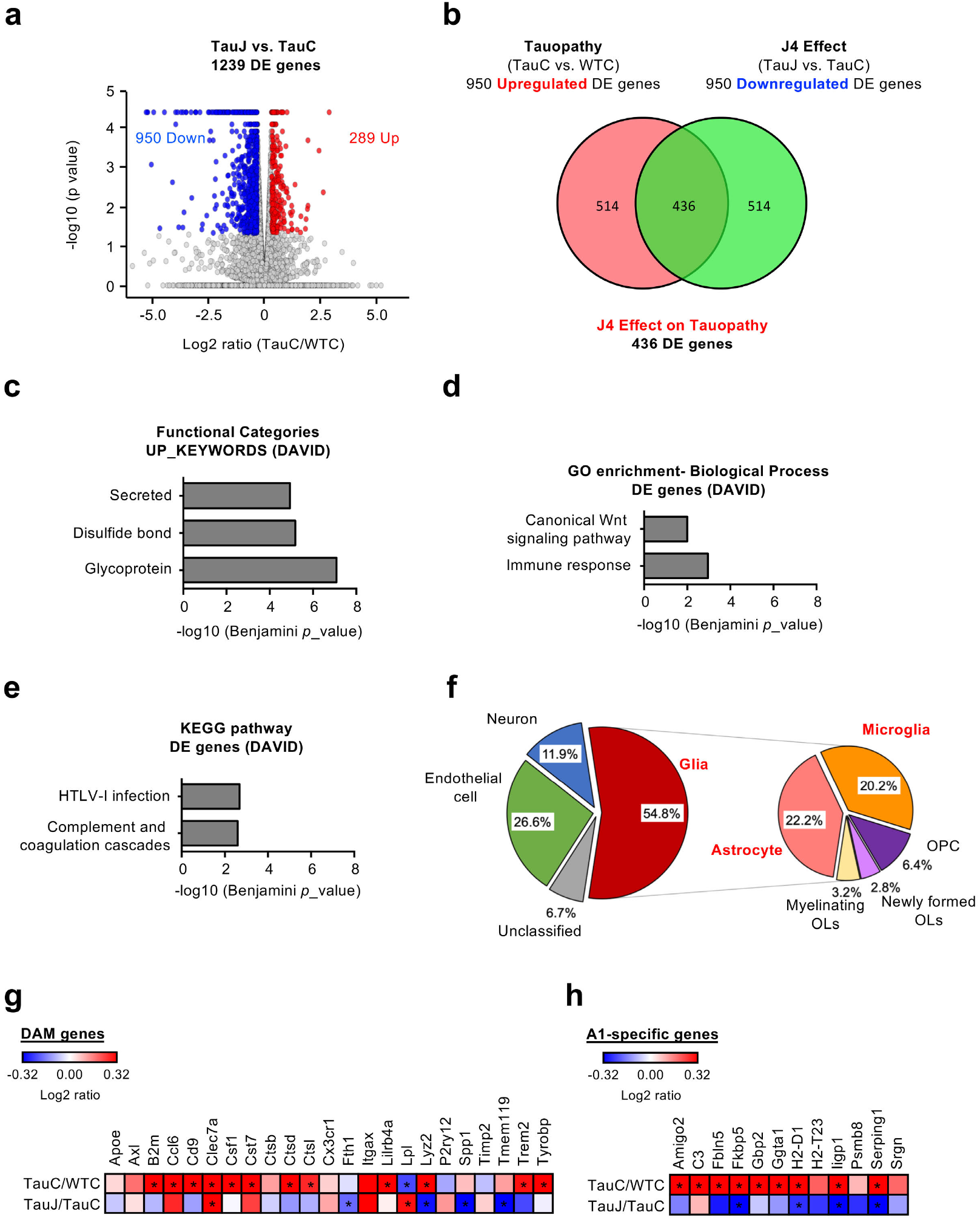
Chronic J4 treatment reduces the immune response and the expression of neurotoxic reactive astrocyte genes in the hippocampi of Tau22 mice. Mice were treated as indicated (control WT mice, WTC; control Tau22 mice, TauC; and J4-treated Tau22 mice, TauJ; n= 3 in each group) from the age of 3-10 months. **(a)** Volcano plot of the DE genes in the TauJ group vs. TauC group. **(b)** Venn diagram showing the 436 overlapping DE genes between the upregulated DE genes in the TauC group vs. WTC group (pink) and the downregulated DE genes in the TauJ group vs. TauC group (green). **(c)** Functional category analysis, **(d)** GO enrichment analysis, and **(e)** KEGG pathway analysis of DE tauopathy-associated genes regulated by J4. **(f)** Pie chart of cell-type-enrichment of 436 tauopathy-associated DE genes regulated by J4. In the volcano plot, the significantly upregulated and downregulated DE genes (absolute log2 ratio ≥ 0.32; *p* < 0.05) are shown in red and blue, respectively. In the Venn diagram and pie chart, the number and percentage of DE genes in each category were shown in each sector. **(g-h)** Heatmap of **(g)** DAM genes and **(h)** A1-specific genes in the TauC/WTC and TauJ/TauC groups. The relative expression level (log2 ratio) of genes is shown on a scale from red (upregulated) to blue (downregulated). Asterisks indicate significant alterations (*p* < 0.05).

### Chronic J4 treatment suppresses the activation of microglia in the hippocampi of Tau22 mice

We next classified the 436 DE genes with expression levels that were normalized by J4 (Fig. 5b) into five types (including neuron-enriched, glia-enriched, endothelial cell-enriched, and unclassified) based on the cell-type information listed in the Brain RNAseq database (https://web.stanford.edu/group/barres_lab/brain_rnaseq.html, (Zhang *et al.*, 2014)). As shown in Fig. 5f, approximately 55% of the DE genes with expression levels that were normalized by J4 were enriched in glial cells, including astrocytes and microglia (Fig. 5f). In the hippocampi of Tau22 mice, enhanced gene signatures for the disease-associated microglia were detected (DAM, (Keren-Shaul *et al.*, 2017); Fig. 5g, Supplementary Table 5). To confirm the activation of microglia, we first determined the number of activated microglia by staining hippocampal sections from Tau22 mice with Iba1 (a marker of microglia) and CD68 (a marker of activated microglia). The intensities of both Iba1 and CD68 were significantly elevated in the hippocampi of TauC mice (11 months old) compared with the hippocampi of WTC mice (Figs. 6a and 6b; \**p* < 0.05, vs WTC mice, one-way ANOVA). We also measured the transcript levels of several factors secreted by activated microglia and known to favor neurotoxic activation of astrocytes (A1 phenotype; (Liddelow *et al.*, 2017)) using RT-qPCR. As shown in Table 1, the levels of *TNF-α* and *C1q (C1qa, C1qb*, and *C1qc*), but not *IL-1α*, were upregulated in the hippocampi of vehicle-treated Tau22 mice compared with the hippocampi of vehicle-treated WT mcie (\**p* < 0.05, versus WTC mice; one-way ANOVA). Immunofluorescence staining and RT-qPCR showed that chronic treatment with J4 prevented the upregulation of *TNF-α*, C1q, and CD68 in Tau22 mice (Figs. 6a-6e, Table 1; ^#^*p* < 0.05, versus TauC mice; one-way ANOVA), suggesting that J4 mitigated microglial inflammation in Tau22 mice.

**Fig. 6.**
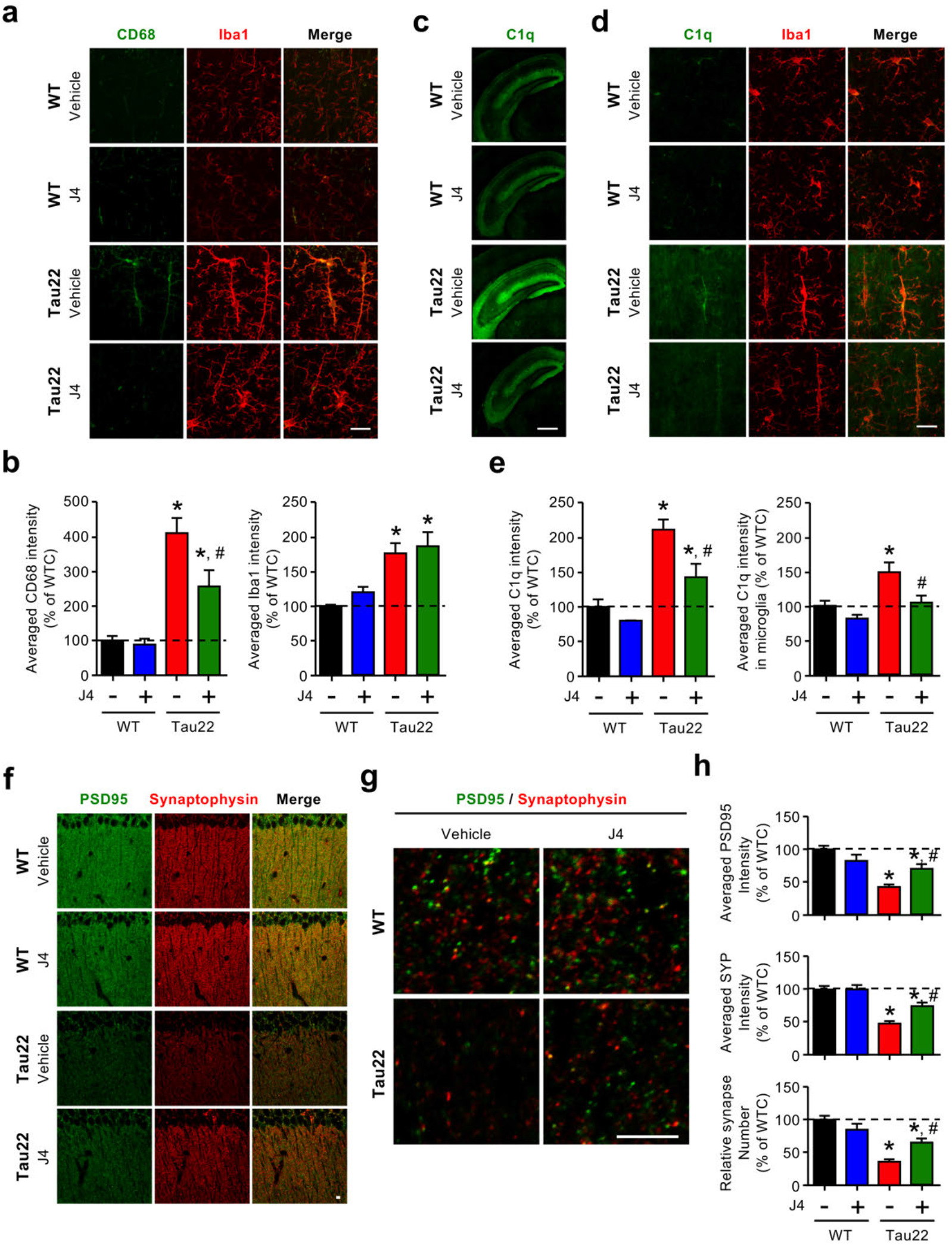
Chronic J4 treatment suppresses the activation of microglia in the hippocampi of Tau22 mice. Mice were treated as indicated (control WT mice, WTC; J4-treated WT mice, WTJ; control Tau22 mice, TauC; and J4-treated Tau22 mice, TauJ; n = 5-7 in each group) from the age of 3-11 months, and their tissues were subjected to IHC staining. The levels of the indicated proteins were evaluated by immunofluorescence staining. **(a-b)** CD68 (green) is a marker of reactive microglia and Iba1 (red) is a marker of microglia, red. (**c-d)**, C1q is shown in green and Iba1 is shown in red. Images were quantified accordingly **(b**, **d)**. Scale bar, 20 μm. The data are expressed as the mean ± S.E.M. \**p* < 0.05 versus the WTC group; ^#^*p* < 0.05 versus the TauC group, one-way ANOVA. **(f-h)** The intensity of synaptic marker expression **(f)** and the number of synapses **(g)** in the hippocampi of treated mice (WTC, black; WTJ, blue; TauC, red; and TauJ, green; n=10-12 in each group) were evaluated by staining with the indicated antibodies (PSD95, green; SYP, red). The quantification results are shown in **(h)**. Scale bar, 5 μm. The data are expressed as the mean ± S.E.M. \**p* < 0.05 versus the WTC group; ^#^*p* < 0.05 versus the TauC group, one-way ANOVA.

**Table 1.**
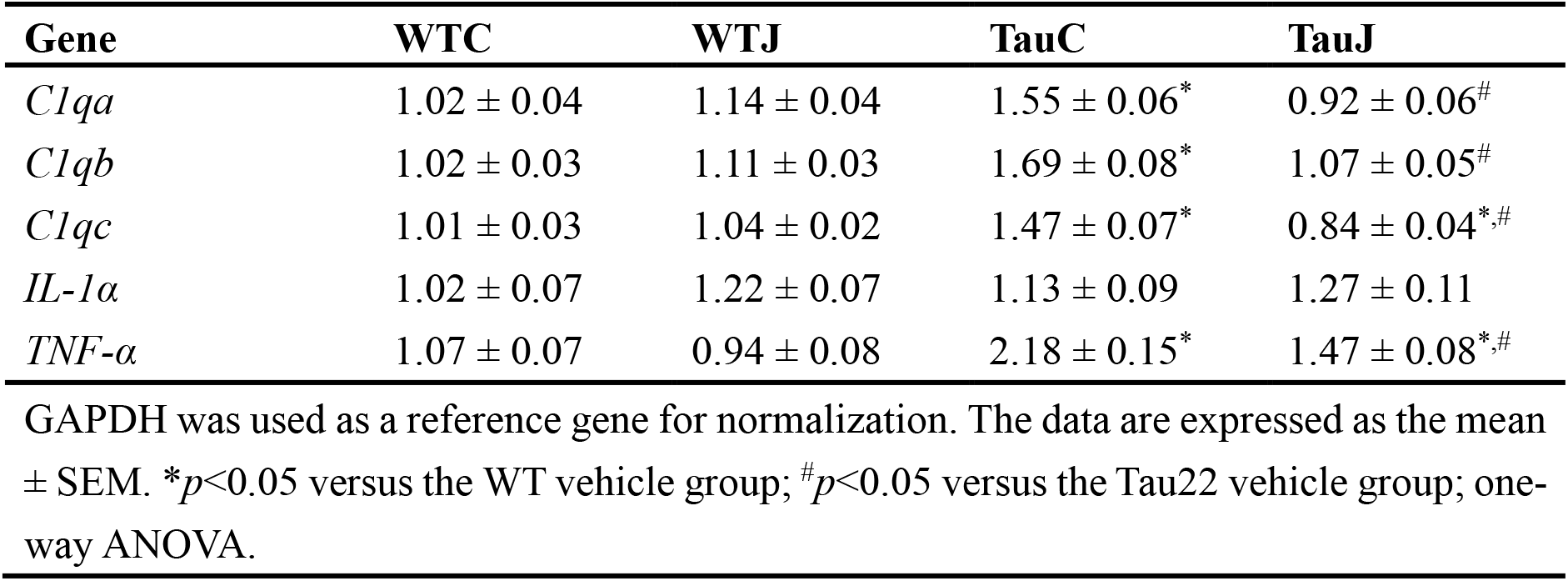
Chronic J4 treatment reduces the transcript levels of activated microglia secreted proinflammatory cytokines in the hippocampi of Tau22 mice. Mice were treated as indicated condition (control WT mice, WTC; J4-treated WT mice, WTJ; control Tau22 mice, TauC; and J4-treated Tau22 mice, TauJ n = 6-9 in each group) from the age of 3-11 months. Gene expression of TNF-α, IL-1α, and C1q (C1qa, C1qb, and C1qc) were analyzed by RT-qPCR. GAPDH was used as a reference gene. The data are expressed as the mean ± S.E.M. *\*p* < 0.05 versus the WTC group; ^#^*p* < 0.05 versus the TauC group, one-way ANOVA.

### Chronic J4 treatment mitigates synaptic loss in Tau22 mice

Earlier studies have demonstrated that C1q is tightly associated with the phagocytic activities of activated microglia for synaptic pruning (Farber *et al.*, 2009), and C1q was recently identified as an important mediator of Tau-induced synaptic loss (Carvalho *et al.*, 2019; Gratuze *et al.*, 2020). We examined the number of synapses by immunofluorescence staining. In the hippocampi of Tau22 mice, the levels of a postsynaptic marker (PSD95) and a presynaptic marker (synaptophysin) were lower than those in the hippocampi of WT mice. Consistent with the rescuing effect of J4 on C1q, chronic J4 treatment restored the levels of PSD95 and synaptophysin in Tau22 mice (Figs. 6f and 6h, \**p* < 0.05, versus WTC mice; ^#^*p* < 0.05, versus TauC mice; one-way ANOVA). We also determined the number of synapses based on the colocalization of PSD95 and synaptophysin. As expected (Chatterjee *et al.*, 2018), the number of synapses in the hippocampi of Tau22 mice was lower than that in the hippocampi of WT mice. In line with the reduction in CD68 and C1q levels by J4 (Table 1, Figs. 6a and 6b), the number of synapses was rescued by chronic J4 treatment in Tau22 mice (Figs. 6g and 6h; \**p* < 0.05, versus WTC mice; ^#^*p* < 0.05, versus TauC mice; one-way ANOVA). This finding is consistent with the alleviation of Tau pathology, synaptic deficits and memory impairment seen in J4-treated Tau22 mice and further supports a detrimental role for microglial activation in tauopathy.

### Chronic J4 treatment suppresses the cytotoxic astrocytes induction in the hippocampi of Tau22 mice

TNF-α and C1q were reported as potent astrocytic activators particularly involved in the establishment of the neurotoxic A1 phenotype (Liddelow *et al.*, 2017). Interestingly, our RNA-seq analysis of the hippocampi of Tau22 mice revealed the upregulation of gene signatures of pan-reactive and cytotoxic A1 astrocytes ((Zamanian *et al.*, 2012; Clarke *et al.*, 2018), Supplementary Table 6 and Fig. 5h). Similarly, in FTD-Tau patients and late stage Alzheimer’s Disease patients (Supplementary Table 7), upregulation of several A1-specific genes (e.g., *GBP2, SERPING1, FKBP5*) was observed. Further, the five A1 astrocyte genes tested were all significantly upregulated in the frontal cortex of FTD-Tau-Pick’s disease patients compared to control subjects, suggesting a link between Tau and astrocyte reactivity (Supplementary Table 7; \**p* < 0.05 versus normal control subjects, two-tailed Student’s *t*-test). Immunofluorescence staining further showed that Tau22 mice (TauC) exhibited higher levels of GFAP (an astrocyte marker) and Lcn2 (a pan reactive astrocyte marker, (Bi *et al.*, 2013)) in the hippocampus than WT mice (WTC; Figs. 7a and 7b; \**p* < 0.05, versus WTC mice, one-way ANOVA), confirming that astrocytes in the hippocampi of Tau22 mice were abnormally activated. The induction of reactive A1 astrocytes appears secondary to microglial activation in the hippocampi of Tau22 mice since activated microglia (CD68-positive) were detected in the hippocampi of young Tau22 mice (4 months old), at which point no reactive astrocytes (Lcn2-positive) were observed (Supplementary Fig. 7).

**Fig. 7.**
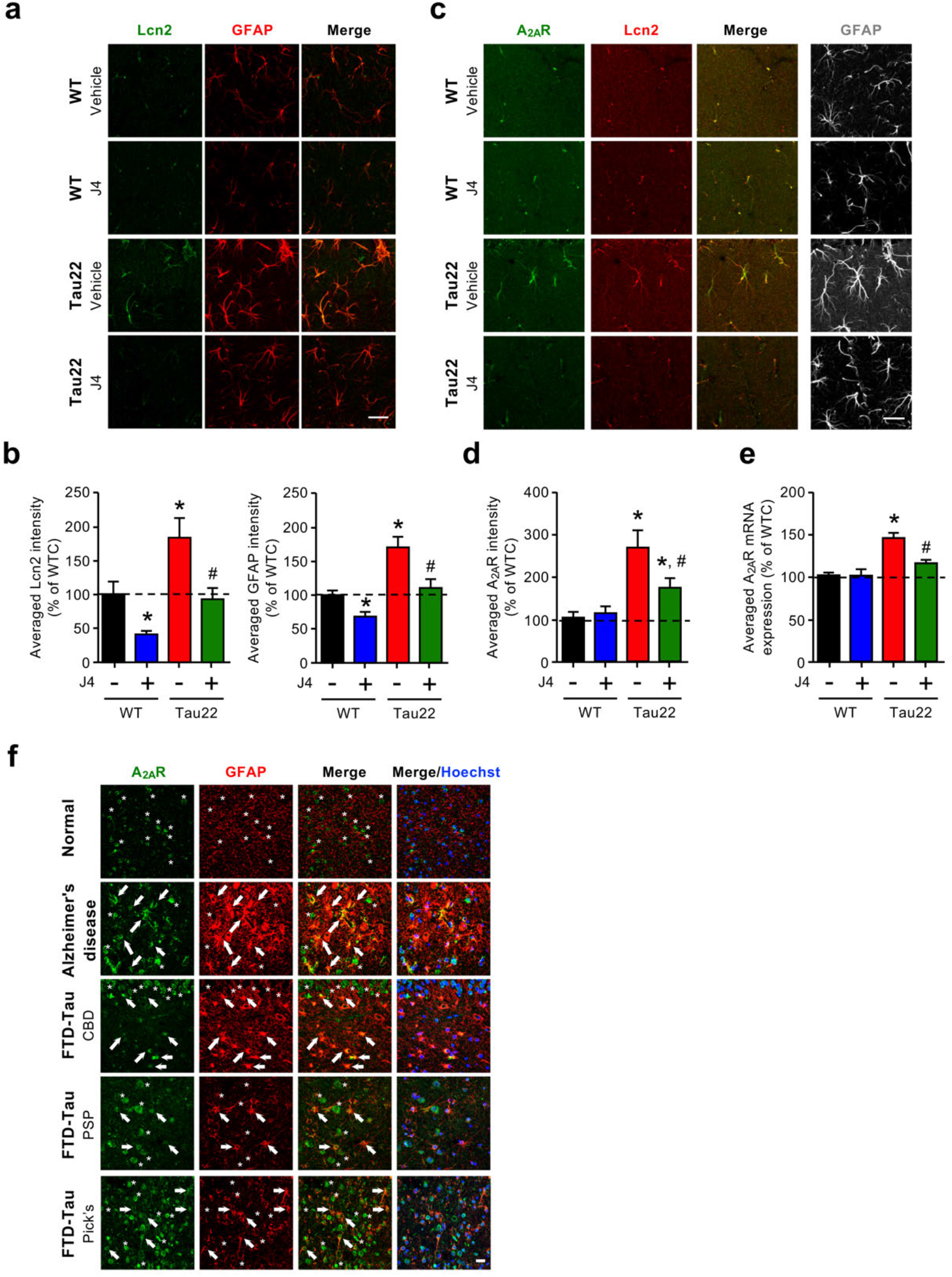
Chronic J4 treatment suppresses the induction of cytotoxic A1 astrocytes in the hippocampi of Tau22 mice. Mice were treated as indicated (control WT mice, WTC; J4-treated WT mice, WTJ; control Tau22 mice, TauC; and J4-treated Tau22 mice, TauJ; n = 5-7 in each group) from the age of 3-11 months. Hippocampal sections (20 μm) were prepared and subjected to IHC staining (a, Lcn2 a marker of reactive astrocytes, green; GFAP a marker of astrocytes, red; c, anti-A_2A_R, green; Lcn2, red; GFAP, grey) and the staining was quantified (b, d). Scale bar, 20 μm. The data are expressed as the mean ± S.E.M. \**p* < 0.05 versus the WTC group; ^#^*p* < 0.05 versus the TauC group, one-way ANOVA. (e) The Adora2a gene expression level in treated mice (WTC, black; WTJ, blue; TauC, red; TauJ, green; n=3-5 in each group) from the age of 3-11 months was examined by RT-qPCR. (f) Posterior hippocampal sections (6 μm) from normal subjects and Alzheimer’s Disease and FTD-Tau (corticobasal degeneration, CBD; progressive supranuclear palsy, PSP; Pick’s disease, Pick’s) patients were subjected to IHC staining. The level of A_2A_R and GFAP were evaluated by staining with the indicated antibodies (human A_2A_R, green; GFAP, red). Scale bar, 20 μm. Arrow, GFAP positive cells; asterisk, GFAP negative cell.

In the Lcn2-positive reactive astrocytes in Tau22 mice, upregulation of the A2A adenosine receptor (A_2A_R) could be detected (Figs. 7c and 7d, \**p* < 0.05, versus WTC mice, one-way ANOVA). The specificity of the anti-A_2A_R antibody was confirmed by negative staining in the brains of A_2A_R-KO mice (Supplementary Fig. 8). In accordance, RT-qPCR analysis showed that the levels of *A_2A_R* transcripts in the hippocampi of Tau22 mice were higher than those in the hippocampi of WT mice (Fig. 7e, \**p* < 0.05, versus WTC mice, one-way ANOVA). An increase in A_2A_R expression in GFAP-positive astrocytes was detected in the brains of postmortem Alzheimer’s Disease and FTLD-Tau patients compared to control subjects (Fig. 7f and Supplementary Fig. 9), further suggesting the link between astrocytic A_2A_R and tauopathies.

According to the reduction of the microglial phenotype as well as *TNFα* and *C1q* expressions (Table 1), chronic J4 treatment normalized not only GFAP and Lcn2 levels but also the pathological upregulation of A1-specific genes and astrocytic A_2A_R expression (Figs. 5h and 7a-d; Supplementary Table 6, ^#^*p* < 0.05, versus TauC mice, one-way ANOVA). Taken together, our data suggest that early Tau-induced microglial activation is likely to promote the activation of neurotoxic astrocytes and pathological upregulation of astrocytic A_2A_R, all blocked by J4.

## Discussion

The present study showed that chronic treatment with J4, an ENT1 blocker, mitigates Tau pathology in Tau22 mice by alleviating not only mitochondrial dysfunction and AMPK overactivation but also the neuroinflammatory status of microglia and astroglia, ultimately attenuating the impairment of compromised synapses as well as spatial learning and memory (Supplementary Fig. 10, Graphic Abstract).

Mitochondrial dysfunction is a major pathogenic feature of Alzheimer’s Disease (Kerr *et al.*, 2017) and is known to facilitate the hyperphosphorylation of Tau, which in turn alters the morphology and functions of mitochondria (DuBoff *et al.*, 2012; Zheng *et al.*, 2020). Therefore, it is not surprising that AMPK, a key energy sensor and an upstream kinase of Tau, is overactivated in the hippocampi of patients with Alzheimer’s Disease or tauopathies (Vingtdeux *et al.*, 2011). One major function of AMPK is the maintenance of cellular energy homeostasis through modulation of the balance between anabolic and catabolic processes (Herzig and Shaw, 2018). In energy-deprived cells (e.g., neurons with impaired mitochondria in the Alzheimer’s Disease brains), a decrease in the ATP/AMP ratio can lead to the activation of AMPK. Accumulating evidence demonstrates that overactivation of AMPK in neurons causes synapse loss via an autophagy-dependent pathway, and links synaptic integrity and energetic failure in neurodegenerative diseases (Domise *et al.*, 2019). In the present study, we found that aberrant AMPK activation was associated with synaptic loss and reduced basal synaptic transmission in the hippocampi of Tau22 mice (Figs. 1e, 3b, 3c, and 6f-6h). The role of AMPK in the regulation of synaptic transmission appears to be complex. When overactivated, AMPK mediates a detrimental effect on neuronal plasticity through inhibition of protein translation via a eukaryotic elongation factor 2 kinase (eEF2K)-dependent pathway Consistently, our findings that chronic treatment with J4 not only suppresses AMPK overactivation but also normalizes impaired neuronal plasticity in both APP/PS1 and Tau22 mice (LTP and LTD, respectively; (Ma *et al.*, 2014); Figs. 1g and 1h).

Treatment with an ENT1 inhibitor (J4) may regulate AMPK by controlling adenosine homeostasis. A recent study demonstrated that genetic deletion of ENT1 reduces adenosine uptake and impacts AMPK activity (Sayama *et al.*, 2020). In the early stages of Alzheimer’s Disease, the level of adenosine is reduced in the frontal cortex but is enhanced in the parietal and temporal cortices. Interestingly, the expression level of adenosine deaminase (ADA) is positively associated with the adenosine level, indicating that ADA may compensate for the loss of adenosine in the human Alzheimer’s Disease brain. In addition, the 5’-nucleotidase activity in different brain areas of Alzheimer’s Disease patients is altered, indicating abnormal purine-related metabolites in the brain (Alonso-Andres *et al.*, 2018). These findings may correlate with our observations of dysregulated mitochondria function and AMPK activity, which is known to disrupt the de novo purine biosynthesis (French *et al.*, 2016; Schmitt *et al.*, 2016), in Tau22 mice (Fig. 3). Additionally, in the symptomatic stage (11 months of age), the expression of 5’-nucleotidase (*CD39* and *CD73*) and *ADA* was significantly enhanced in the hippocampi of Tau22 mice compared to the hippocampi of control mice (Supplementary Table 8). These data showed that purine metabolism in the hippocampi of Tau22 mice is abnormal. Chronic J4 treatment rescued the altered expression of several enzymes involved in adenosine metabolism (Supplementary Table 8), suggesting that J4 normalized aberrant purine metabolism in the Alzheimer’s Disease brain. Future analyses of local concentrations of adenosine and ATP using cell typespecific reporters are warranted.

Although pathogenic Tau is specifically expressed in the neurons of Tau22 mice, reactive microglia and astrocytes have been found near neurons that contain high levels of pathogenic Tau (Schindowski *et al.*, 2006), suggesting that degenerating neurons may trigger abnormal gliosis. In another mouse model of tauopathy (rTg4510), RNA-seq analysis revealed that microglia respond to stress from neurons expressing pathogenic tau^P301L^, and alter their gene expression profiles (Wang *et al.*, 2018). In the present study, our RNA-seq analysis of Tau22 mice showed upregulation of several genes associated with the disease-associated microglia (DAM, (Keren-Shaul *et al.*, 2017)) in mice and patients with Alzheimer’s Disease (Fig. 5g, and Supplementary Table 5). Notably, J4 treatment not only ameliorated the energy dysfunction and Tau pathology in neurons (Figs. 1–3) and markedly reduced microglia activation and rescued the synapse loss associated with the phagocytic activities of activated microglia (Fig. 6) (Dejanovic *et al.*, 2018; Carvalho *et al.*, 2019; Gratuze *et al.*, 2020). We also performed RT-qPCR and established that microglial factors known to promote neurotoxic activation of astrocytes were upregulated in Tau22 mice (*C1q* and *TNFa;* Table 1, Fig. 6). In line with the suppression of microglial activation, J4 treatment also significantly reduced astrocytic activation, with a particular impact on the A1 signature (Fig. 7). Here, we show, for the first time, a pathological activation of the A1 neurotoxic astrocytic phenotype in a tauopathy context using both human and mouse samples (Supplementary Tables 6 and 7), which may be related to the production of A1-promoting factors by microglia. Our RNA-seq analysis of Tau22 mice also emphasizes the upregulation of a disease-associated astrocytes (DAAs) signature, recently detected in an amyloid Alzheimer’s Disease mouse model (5XFAD) (Habib *et al.*, 2020) (Supplementary Table 9). Together, our observations are of particular importance since glial activation has been shown to be instrumental in Alzheimer’s Disease (Laurent *et al.*, 2018). More recently, astrogliosis has been also involved (Balu *et al.*, 2019). For instance, suppression of the Stat3-mediated astrogliosis improved behavioral impairments in APP/PS1 mice (Reichenbach *et al.*, 2019). In accordance, the beneficial effects of J4 on memory alterations and neuroplasticity (Figs. 1–3) is associated with the normalization of both A1-promoting factors and A1 astrocytes in Tau22 mice (Supplementary Table 6). Treatment with J4 normalized 60% of the elevated DAA genes in Tau22 mice (Supplementary Table 9), further supporting the beneficial effect of J4 on the prevention of abnormal astrocytic activation.

As detailed in our previous study (Lee *et al.*, 2018), J4 has a very small or no effect on the major adenosine receptors in the brain (A_1_R and A_2A_R). A direct contribution of adenosine receptors to the beneficial effect of J4 appears therefore unlikely. Surprisingly, we found that the expression of astroglial A_2A_R was significantly increased not only in the hippocampi of patients with tauopathies but also in Tau22 mice (Fig. 7). The function of astrocytic A_2A_R remains elusive. However, it has been associated with impaired memory in hAPP mice (Orr *et al.*, 2015). A recent study also demonstrated that overactivation of A_2A_R in astrocytes markedly alters transcriptional changes, particularly affecting immune response (Paiva *et al.*, 2019). Interestingly, activation of A_2A_R in astrocytes enhances glutamate release and inhibits glutamate uptake (Matos *et al.*, 2013; Cervetto *et al.*, 2017). An increase in astrocytic A_2A_R expression may therefore be prone to increase the level of extracellular glutamate and facilitate excitotoxicity. The present study indicated that, while it does not directly act on A_2A_Rs, J4 leads to a significant reduction in astrocytic receptor levels in Tau22 mice, suggesting that a reduction in astrocytic A_2A_R contributes to improving the phenotype, which is in line with a previous work showing that global A_2A_R blockade phenocopies the effect of J4 in the same Tau22 line by reducing astroglial activation (Laurent *et al.*, 2016).

## Conclusion

In summary, we provide evidence that an ENT1 inhibitor (J4) rescues the energy dysfunction (including mitochondrial impairment and AMPK overactivation) and pathological glial activation, and subsequently improves synaptic function and memory in tauopathy. Modulation of adenosine homeostasis by an ENT1 inhibitor (J4) therefore deserves further development in tauopathies and Alzheimer’s Disease.

## Acknowledgements

We are grateful to the Proteomics Core Facility of the Institute of Biomedical Sciences, Academia Sinica and the Neuroscience core facility, Academia Sinica for the LC/MS/MS analysis and the electrophysiology analysis. We acknowledge the UC Davis Alzheimer’s Disease Center (USA) for providing human brain specimens from normal, Alzheimer’s Disease, and FTD-Tau subjects. We also thank Dr. Peter Davies (The Feinstein Institute for Medical Research, New York, USA) for generously providing disease-specific conformational modified tau antibody (MC1).

## Supplementary material

Supplementary material is available at Brain online.

