## Supplementary material for "A novel equilibrative nucleoside transporter 1 inhibitor alleviates Tau-mediated neurodegeneration"

#### Inventory of Supplemental Information

a) Supplemental Tables

b) Supplemental Figures and Legends

c) Supplemental Experimental Procedures

d) Supplemental References

### Supplemental Tables

**Supplementary Table 1. Human cases analyzed in this study**

| DX | NA ID | Braak | Age | Sex | PMI (hrs) | Region | Application |
|---|---|---|---|---|---|---|---|
| Alzheimer's disease | 4543 | 6 | 71 | M | 9 | Hp | IF |
| Alzheimer's disease | 4552 | 6 | 64 | F | 6 | Hp | IF |
| Alzheimer's disease | 4580 | 6 | 73 | M | 12 | Hp | IF |
| Alzheimer's disease | 4851 | 6 | 69 | M | 17 | Hp | IF |
| Alzheimer's disease | 4872 | NA | 71 | F | NA | Hp | IF |
| Alzheimer's disease | 4966 | 6 | 75 | F | 17 | Hp | IF |
| FTD-Tau (CBD) | 4855 | 5 | 72 | F | 19.5 | Hp | IF |
| FTD-Tau (Pick's) | 4439 | 0 | 69 | M | 12 | Hp | IF |
| FTD-Tau (Pick's) | 4798 | 0 | 74 | F | 18 | Hp | IF |
| FTD-Tau (Pick's) | 4814 | 0 | 68 | F | 24 | Hp | IF |
| FTD-Tau (Pick's) | 4933 | 2 | 74 | M | 19 | Hp | IF |
| FTD-Tau (PSP) | 4744 | 1 | 65 | F | 8 | Hp | IF |
| Normal | 4706 | 1 | 75 | F | 3.5 | Hp | IF |
| Normal | 4727 | 1 | 78 | F | NA | Hp | IF |
| Normal | 4768 | 1 | 82 | F | 14.17 | Hp | IF |
| Normal | 4795 | 2 | 75 | M | 18 | Hp | IF |
| Normal | 4858 | 2 | 78 | F | 40.83 | Hp | IF |
| Normal | 4894 | 2 | 75 | M | 49.5 | Hp | IF |
| Alzheimer's disease /MCI | C09-19424 | 4 | 88 | F | 16 | BA10 | RT-qPCR |
| Alzheimer's disease /MCI | G 56 | 4 | 90 | F | 4 | BA10 | RT-qPCR |
| Alzheimer's disease /MCI | G 73 | 3 | 92 | F | 4 | BA10 | RT-qPCR |
| Alzheimer's disease /MCI | G 89 | 4 | 68 | M | 6 | BA10 | RT-qPCR |
| Alzheimer's disease /MCI | P 7024 | 4 | 76 | F | 28 | BA10 | RT-qPCR |
| Alzheimer's disease | C07-20298 | 6 | 77 | F | 15 | BA10 | RT-qPCR |
| Alzheimer's disease | C07-20991 | 6 | 80 | F | 5 | BA10 | RT-qPCR |
| Alzheimer's disease | C08-03086 | 6 | 86 | M | 10 | BA10 | RT-qPCR |
| Alzheimer's disease | C08-10048 | 5 | 62 | M | 10 | BA10 | RT-qPCR |
| Alzheimer's disease | C08-28113 | 6 | 86 | F | 8 | BA10 | RT-qPCR |

| | | | | | | | |
|---|---|---|---|---|---|---|---|
| Alzheimer’s disease | C08-31992 | 5 | 89 | F | 24 | BA10 | RT-qPCR |
| Alzheimer’s disease | C09-34184 | 6 | 73 | F | 22 | BA10 | RT-qPCR |
| Alzheimer’s disease | C10-03400 | 6 | 72 | F | 5.5 | BA10 | RT-qPCR |
| Alzheimer’s disease | C11-34515 | 6 | 67 | M | 9.5 | BA10 | RT-qPCR |
| Alzheimer’s disease | C11-36501 | 6 | 65 | M | 36 | BA10 | RT-qPCR |
| Alzheimer’s disease | C12-00612 | 6 | 61 | F | 17 | BA10 | RT-qPCR |
| Alzheimer’s disease | C12-05247 | 6 | 83 | F | 3 | BA10 | RT-qPCR |
| Alzheimer’s disease | C12-10400 | 5 | 89 | H | 20 | BA10 | RT-qPCR |
| Alzheimer’s disease | G 109 | 5 | 92 | F | NA | BA10 | RT-qPCR |
| FTD-Tau (CBD) | C08-09824 | NA | 75 | F | 7 | BA10 | RT-qPCR |
| FTD-Tau (CBD) | C08-18677 | NA | 76 | M | 19 | BA10 | RT-qPCR |
| FTD-Tau (CBD) | C08-29472 | NA | 71 | M | 5 | BA10 | RT-qPCR |
| FTD-Tau (CBD) | P 5925 | NA | 72 | F | 15 | BA10 | RT-qPCR |
| FTD-Tau (CBD) | P 8671 | NA | 81 | F | 33 | BA10 | RT-qPCR |
| FTD-Tau (PSP) | C08-04599 | NA | 77 | F | 17 | BA10 | RT-qPCR |
| FTD-Tau (PSP) | C08-18602 | 3 | 75 | M | 7 | BA10 | RT-qPCR |
| FTD-Tau (PSP) | C09-17736 | NA | 65 | M | 18 | BA10 | RT-qPCR |
| FTD-Tau (PSP) | C10-06503 | NA | 82 | F | 11 | BA10 | RT-qPCR |
| FTD-Tau (PSP) | C10-07949 | NA | 77 | F | 24 | BA10 | RT-qPCR |
| FTD-Tau (PSP) | C12-37132 | NA | 77 | F | 16 | BA10 | RT-qPCR |
| FTD-Tau (PSP) | C14-00229 | NA | 89 | M | 36 | BA10 | RT-qPCR |
| FTD-Tau (PSP) | P 3814 | NA | 76 | M | 7 | BA10 | RT-qPCR |
| FTD-Tau (PSP) | P 4117 | NA | 70 | M | NA | BA10 | RT-qPCR |
| FTD-Tau (PSP) | P 7622 | NA | 80 | F | 22 | BA10 | RT-qPCR |
| FTD-Tau (Pick's) | C09-21706 | NA | 71 | M | 21 | BA10 | RT-qPCR |
| FTD-Tau (Pick's) | C10-05402 | NA | 68 | F | 11 | BA10 | RT-qPCR |
| FTD-Tau (Pick's) | C11-33155 | NA | 68 | M | 15 | BA10 | RT-qPCR |
| FTD-Tau (Pick's) | P 3538 | NA | 60 | M | NA | BA10 | RT-qPCR |
| FTD-Tau (Pick's) | P 8245 | NA | 67 | F | NA | BA10 | RT-qPCR |
| Normal | C06-14473 | 2 | 72 | M | 18 | BA10 | RT-qPCR |
| Normal | C08-05295 | 0 | 52 | F | 28 | BA10 | RT-qPCR |
| Normal | P 3549 | 2 | 69 | M | 6 | BA10 | RT-qPCR |
| Normal | P 3862 | 1-2 | 58 | M | 5.5 | BA10 | RT-qPCR |
| Normal | P 4078 | 2 | 73 | M | 10 | BA10 | RT-qPCR |
| Normal | P 8730 | 1-2 | 82 | M | NA | BA10 | RT-qPCR |
| Normal | P 8866 | 1-2 | 84 | M | 15.5 | BA10 | RT-qPCR |
| Normal | G 15 | 1 | 80 | M | 26 | BA10 | RT-qPCR |
| Normal | G 18 | 1 | 76 | F | 8 | BA10 | RT-qPCR |

| | | | | | | | |
|---|---|---|---|---|---|---|---|
| Normal | G 69 | 1 | 95 | F | 3 | BA10 | RT-qPCR |
| FTD-Tau (P301L) | A02 1102 | NA | 43 | M | 6 | TpCx | RT-qPCR |
| FTD-Tau (P301L) | A02 1213 | NA | 66 | M | 30 | TpCx | RT-qPCR |
| FTD-Tau (P301L) | A06 2025 | NA | 65 | F | NA | TpCx | RT-qPCR |
| Normal | A09 0606 | NA | 70 | M | 30 | TpCx | RT-qPCR |
| Normal | A11 1620 | NA | 72 | M | 7 | TpCx | RT-qPCR |
| Normal | A14 0019 | NA | 53 | F | 29 | TpCx | RT-qPCR |

Alzheimer's disease/MCI, Mild cognitive impairment with early Alzheimer's disease diagnosis; FTD, Frontotemporal dementia; CBD, Corticobasal degeneration; PSP, Progressive supranuclear palsy; PMI, Postmortem interval; M, Male; F, Female; Hp, Posterior hippocampus; BA10, Prefrontal cortex Brodmann area 10; TpCx, Temporal Cortex; NA, not available; IF, immunofluorescence; RT-qPCR, quantitative reverse transcription PCR.

**Supplementary Table 2. Sequences of the RT-qPCR primers used in this study**

| Gene name | Primer sequence | | Product size (bp) |
|---|---|---|---|
| mAda | Forward | 5'-AACATTATCGGCATGGACAAGC-3' | 132 bp |
| | Reverse | 5'-TGCCTTCATCTCCACAAACTCG-3' | |
| mAdk | Forward | 5'-GTGCTATTTGGAATGGGGAAT-3' | 207 bp |
| | Reverse | 5'-CAACCACTGAGCCACTTTCAT-3' | |
| mAdora2a | Forward | 5'-CTGCTTTGTCCTGGTCCTCAC-3' | 114 bp |
| | Reverse | 5'-ATACCCGTCACCAAGCCATT-3' | |
| mAdora1 | Forward | 5'-TTGCTATTGCTGTGGATCGGT-3' | 146 bp |
| | Reverse | 5'-GTTCCAGCCAAACATGGGTGT-3' | |
| mC1qa | Forward | 5'-GAGCACCCAACGGGAAGGAT-3' | 111 bp |
| | Reverse | 5'-GGATACCAGTCCGGATGCCA-3' | |
| mC1qb | Forward | 5'-CACAGAACACCAGGATTCCATAC-3' | 109 bp |
| | Reverse | 5'-GGAGAAAACCTAGAAGCAGCAGT-3' | |
| mC1qc | Forward | 5'-GCGATGAGGTGTGGCTATCA-3' | 174 bp |
| | Reverse | 5'-GGAAGAGGTCTGAGTGAGGATG-3' | |
| mCD39 (Entpd1) | Forward | 5'-GTACCTGAGTGAGTACTGCTTCTC-3' | 165 bp |
| | Reverse | 5'-GCTGGGATGTTGGTCAAGTTC-3' | |
| mCD73 (Nt5e) | Forward | 5'-CAAATCCCACACAACCACTG-3' | 158 bp |
| | Reverse | 5'-TGCTCACTTGGTCACAGGAC-3' | |
| mENT1 | Forward | 5'-ATCAATCATGCGAAAGCA-3' | 131 bp |
| | Reverse | 5'-GCAGGTGAAGACCAGCAA-3' | |
| mENT2 | Forward | 5'-CATGGAAACTGAGGGGAAGA-3' | 186 bp |
| | Reverse | 5'-GTTCCAAAGGCCTCACAGAG-3' | |
| mENT3 | Forward | 5'-AAACTCCGAAACTGCTCC-3' | 130 bp |
| | Reverse | 5'-GGAAGTTAGCCACCAGGA-3' | |
| mENT4 | Forward | 5'-GCGGCTGTGCTCCTAAAC-3' | 148 bp |
| | Reverse | 5'-CGTAAGCCTGGTCGTGAG-3' | |
| mGAPDH | Forward | 5'-TGACATCAAGAAGGTGGTGAAG-3' | 109 bp |
| | Reverse | 5'-AGAGTGGGAGTTGCTGTTGAAG-3' | |
| mIL1-α | Forward | 5'-TGCAGTCCATAACCCATGATC-3' | 113 bp |
| | Reverse | 5'-GACAAACTTCTGCCTGACGAG-3' | |
| mTNFα | Forward | 5'-TCTTCTCATTCCTGCTTGTGG-3' | 130 bp |
| | Reverse | 5'-AGGGTCTGGGCCATAGAACT-3' | |

The sequences of the forward and reverse primers (5'→3' direction) for each gene of interest used for RT-qPCR assays are listed.

**Supplementary Table 3. The list of TaqMan probes used in this study**

| Assay ID | Gene | Target species | Assay design | Amplicon length (bp) |
|---|---|---|---|---|
| Hs00163811_m1 | C3 | Human | Probe spans exons | 88 bp |
| Hs00163781_m1 | SERPING1 | Human | Probe spans exons | 70 bp |
| Hs01561006_m1 | FKBP5 | Human | Probe spans exons | 75 bp |
| Hs01056636_m1 | FBLN5 | Human | Probe spans exons | 65 bp |
| Hs00894837_m1 | GBP2 | Human | Probe spans exons | 84 bp |
| Hs99999903_m1 | ACTB | Human | Amplicon spans exons | 171 bp |

The TaqMan assay ID and design for each gene of interest used for RT-qPCR assays are listed.

#### Supplementary Table 4. List of antibodies

| Name | Abbreviation | Source | Species reactivity | Dilution | Source (Cat No.) |
|---|---|---|---|---|---|
| Anti-Glial fibrillary acidic protein | GFAP | Rabbit | Mouse, Human | IF: 1:500 | Sigma-Aldrich (G9269) |
| Anti-Glial fibrillary acidic protein | GFAP | Mouse | Mouse, Human | IF: 1:200 | Sigma-Aldrich (G3893) |
| Anti-Iba1 | Iba1 | Rabbit | Mouse, Human | IF: 1:500 | Wako (019-19741) |
| AIF-1/Iba1 | Iba1 | Goat | Mouse, Human | IF: 1:300 | Novus Biologicals (NB100-1028) |
| Lipocalin-2/ NGAL | Lcn2 | Goat | Mouse | IF: 1:200 | R&D (AF1857) |
| Anti-CD68 (ED1) | CD68 | Mouse | Mouse, Human | IF: 1:100 | Abcam (ab31630) |
| Phospho-Tau (Ser199) | p-Tau$^{Ser199}$ | Rabbit | Mouse, Human | WB: 1:1000 | Thermo Fisher Scientific (44-734G) |
| Phospho-Tau (Thr212, Ser214) | AT100 | Mouse | Human | IF: 1:1000<br>WB: 1:1000 | Thermo Fisher Scientific (MN1060) |
| Phospho-Tau (Ser202, Thr205) | AT8 | Mouse | Mouse, Human | IF: 1:2000 | Thermo Fisher Scientific (MN1020) |
| Phospho-Tau (Ser262) | p-Tau$^{Ser262}$ | Rabbit | Mouse, Human | WB: 1:1000 | Thermo Fisher Scientific (44-750G) |
| Phospho-Tau (Ser396) | p-Tau$^{Ser396}$ | Rabbit | Mouse, Human | WB: 1:2000 | Thermo Fisher Scientific (44-752G) |
| Phospho-Tau (Ser422) | p-Tau$^{Ser422}$ | Rabbit | Mouse, Human | WB: 1:1000 | Thermo Fisher Scientific (44-764G) |
| Phospho-Tau (Thr181) | AT270 | Mouse | Human | WB: 1:1000 | Thermo Fisher Scientific (MN1050) |
| Tau (MC1) | MC1 | Mouse | Mouse, Human | IF: 1:1000<br>WB: 1:1000 | Gift from Dr. Peter Davies |
| Phospho-Tau (Ser396, Ser404) (PHF1) | PHF1 | Mouse | Human | WB: 1:1000 | Gift from Dr. Peter Davies |
| Anti-Tau | Tau-5 | Mouse | Mouse, Human | WB: 1:2500 | Thermo Fisher Scientific (AHB0042) |
| Anti-Human Tau | HT7 | Mouse | Human | WB: 1:2000 | Thermo Fisher Scientific (MN1000) |
| Phospho-AMPK□ (Thr172) | p-AMPK$^{Thr172}$ | Rabbit | Mouse, Human | IF: 1:50 | Cell Signaling (#2535) |
| Anti-ATP5A antibody [15H4C4] | ATP5A | Mouse | Mouse, Human | IF: 1:200 | Abcam (ab14748) |
| Anti-C1q antibody [4.8] | C1q | Rabbit | Mouse | IF: 750 | Abcam (ab227072) |
| Anti-PSD95, clone 7E3-1B8 | PSD95 | Mouse | Mouse, Human | IF: 1:250 | Millipore (MAB1598) |
| SYP (C-20) | SYP | Goat | Mouse, Human | IF: 1:100 | Santa Cruz (Sc-7568) |
| Anti-$A_{2A}R$ | $A_{2A}R$ | Guinea pig | Mouse | IF: 200 | Frontier Institute (ABN454) |
| Anti-$A_{2A}R$ | $A_{2A}R$ | Rabbit | Human | IF: 1:100 | Homemade |

**Supplementary Table 5. Disease-associated microglia (DAM) genes**

| Type | Gene name | TauC/WTC | *p*_value | TauJ/TauC | *p*_value |
|---|---|---|---|---|---|
| DAM genes | *Apoe* | 0.05 | 0.62 | -0.07 | 0.54 |
| | *Axl* | 0.18 | 0.10 | 0.04 | 0.73 |
| | *Ccl6* | **0.69*** | 0.00 | 0.27 | 0.14 |
| | *Cd9* | **0.58*** | 0.00 | -0.12 | 0.15 |
| | *Clec7a* | **2.91*** | 0.00 | **0.68*** | 0.01 |
| | *Csf1* | **0.37*** | 0.00 | -0.01 | 0.89 |
| | *Cst7* | **2.80*** | 0.00 | 0.24 | 0.31 |
| | *Ctsb* | 0.12 | 0.14 | -0.06 | 0.44 |
| | *Ctsd* | **0.41*** | 0.00 | -0.13 | 0.10 |
| | *Ctsl* | **0.27*** | 0.00 | -0.10 | 0.29 |
| | *Cx3cr1* | 0.06 | 0.69 | 0.11 | 0.45 |
| | *Fth1* | -0.04 | 0.63 | **-0.19**# | 0.03 |
| | *Itgax* | 3.24 | 1.00 | 0.34 | 0.37 |
| | *Lilrb4a* | **1.54*** | 0.00 | 0.02 | 0.96 |
| | *Lpl* | **-0.23*** | 0.03 | **0.30*** | 0.01 |
| | *Lyz2* | **1.06*** | 0.00 | **-0.30**# | 0.01 |
| | *P2ry12* | -0.11 | 0.43 | 0.13 | 0.38 |
| | *Spp1* | 0.08 | 0.58 | **-0.49**# | 0.00 |
| | *Timp2* | -0.08 | 0.30 | 0.06 | 0.42 |
| | *Tmem119* | 0.12 | 0.21 | **-0.32**# | 0.00 |
| | *Trem2* | **0.30*** | 0.04 | -0.23 | 0.15 |
| | *Tyrobp* | **0.33*** | 0.02 | -0.02 | 0.90 |

Mice were treated as indicated conditions (control WT mice, WTC; control Tau22 mice, TauC; and J4-treated Tau22 mice, TauJ; n= 3 in each group) from the age of 3-10 months. The hippocampus was harvested carefully and subjected to RNA-seq analysis. The relative levels (log2 ratio) of DAM genes are listed. *$p<0.05$ versus WTC mice; #$p<0.05$ versus TauC mice.

**Supplementary Table 6. PAN-reactive and A1-specific genes**

| Type | Gene name | TauC/WTC | *p*_value | TauJ/TauC | *p*_value |
|---|---|---|---|---|---|
| **PAN-reactive genes** | *Aspg* | **0.81*** | 0.01 | -0.25 | 0.40 |
| | *Cd44* | 0.27 | 0.08 | **0.31#** | 0.04 |
| | *Cp* | 0.28 | 0.09 | -0.18 | 0.24 |
| | *Cxcl10* | 0.60 | 0.29 | 0.48 | 0.37 |
| | *Gfap* | **0.73*** | 0.00 | -0.14 | 0.22 |
| | *Hsbp1* | -0.12 | 0.14 | -0.04 | 0.60 |
| | *Lcn2* | 3.44 | 0.28 | -2.40 | 0.15 |
| | *Osmr* | **0.56*** | 0.00 | 0.00 | 0.98 |
| | *S1pr3* | 0.07 | 0.59 | 0.02 | 0.89 |
| | *Serpina3n* | **0.52*** | 0.00 | 0.03 | 0.72 |
| | *Steap4* | **1.58*** | 0.00 | **-1.53#** | 0.00 |
| | *Timp1* | 1.16 | 1.00 | -0.12 | 1.00 |
| | *Vim* | **0.37*** | 0.00 | **-0.22#** | 0.03 |
| **A1-specific genes** | *Amigo2* | **0.63*** | 0.00 | -0.16 | 0.10 |
| | *C3* | **1.33*** | 0.00 | 0.09 | 0.83 |
| | *Fbln5* | **0.61*** | 0.00 | -0.29 | 0.11 |
| | *Fkbp5* | **0.42*** | 0.00 | **-0.38#** | 0.00 |
| | *Gbp2* | **0.84*** | 0.00 | -0.07 | 0.66 |
| | *Ggta1* | **0.68*** | 0.00 | -0.13 | 0.40 |
| | *H2-D1* | **0.56*** | 0.00 | **-0.26#** | 0.01 |
| | *H2-T23* | 0.20 | 0.30 | -0.25 | 0.18 |
| | *Iigp1* | **0.85*** | 0.00 | **-0.41#** | 0.02 |
| | *Psmb8* | 0.09 | 0.74 | -0.26 | 0.35 |
| | *Serping1* | **0.46*** | 0.00 | **-0.49#** | 0.00 |
| | *Srgn* | 0.19 | 0.16 | -0.12 | 0.38 |

Mice were treated as indicated conditions (control WT mice, WTC; control Tau22 mice, TauC; and J4-treated Tau22 mice, TauJ; n= 3 in each group) from the age of 3-10 months. The hippocampus was harvested carefully and subjected to RNA-seq analysis. The data are expressed as the log2 ratio. *$p$<0.05 versus the WT vehicle group; $^{\#}p$<0.05 versus the Tau22 vehicle group.

**Supplementary Table 7. The expression of the A1-specific genes in the frontal cortex of AD and FTD-Tau patients**

| Gene name | Group | *C3* | *FBLN5* | *FKBP5* | *GBP2* | *SERPING1* |
|---|---|---|---|---|---|---|
| **Alzheimer's disease/MCI** | Normal (n=10) | 1.00 ± 0.16 | 1.00 ± 0.15 | 1.00 ± 0.21 | 1.00 ± 0.19 | 1.00 ± 0.16 |
| | AD/MCI (n=5) | 1.07 ± 0.59 | 0.49 ± 0.35 | 1.31 ± 0.47 | 1.26 ± 1.11 | 1.05 ± 0.46 |
| **Alzheimer's disease** | Normal (n=10) | 1.00 ± 0.16 | 1.00 ± 0.15 | 1.00 ± 0.21 | 1.00 ± 0.19 | 1.00 ± 0.16 |
| | AD (n=14) | 1.16 ± 0.16 | 0.93 ± 0.40 | 1.77 ± 0.25* | 1.18 ± 0.34 | 1.19 ± 0.15 |
| **FTD-Tau-CBD** | Normal (n=10) | 1.00 ± 0.16 | 1.00 ± 0.15 | 1.00 ± 0.21 | 1.00 ± 0.19 | 1.00 ± 0.16 |
| | CBD (n=5) | 1.77 ± 0.67 | 1.11 ± 0.21 | 4.45 ± 1.46* | 5.28 ± 1.05* | 2.20 ± 0.49* |
| **FTD-Tau-P301L** | Normal (n=3) | 1.00 ± 0.56 | 1.00 ± 0.25 | 1.00 ± 0.27 | 1.00 ± 0.27 | 1.00 ± 0.37 |
| | P301L (n=3) | 6.27 ± 4.34 | 2.31 ± 0.80 | 2.76 ± 0.71 | 2.80 ± 0.50* | 2.71 ± 1.25 |
| **FTD-Tau-Pick** | Normal (n=10) | 1.00 ± 0.16 | 1.00 ± 0.15 | 1.00 ± 0.21 | 1.00 ± 0.19 | 1.00 ± 0.16 |
| | Pick's (n=5) | 2.31 ± 0.42* | 3.62 ± 1.36* | 2.96 ± 0.76* | 6.04 ± 0.99* | 3.12 ± 0.49* |
| **FTD-Tau-PSP** | Normal (n=10) | 1.00 ± 0.16 | 1.00 ± 0.15 | 1.00 ± 0.21 | 1.00 ± 0.19 | 1.00 ± 0.16 |
| | PSP (n=10) | 0.94 ± 0.28 | 1.01 ± 0.07 | 1.76 ± 0.34 | 2.06 ± 0.46* | 1.53 ± 0.20 |

The expression levels of A1-specific genes in the frontal cortices of AD (AD/MCI, Braak 3-4, n=5; AD, Braak 5-6, n=14) and FTD-Tau patients (CBD, n=5; P301L, n=3; Pick, n=5; PSP, n=10) and normal control subjects (n=3-10) were analyzed by RT-qPCR. The relative expression level of genes is listed. ACTB was used as a reference gene for normalization. The data are expressed as the mean ± SEM. *$p$<0.05 versus the normal control subjects; two-tailed Student's $t$-test. AD, Alzheimer's disease; FTD, Frontotemporal dementia; FTLD, Frontotemporal lobar degeneration; CBD, Corticobasal degeneration; PSP, Progressive supranuclear palsy.

**Supplementary Table 8. Enzymes involved in adenosine homeostasis**

| Gene | WTC | WTJ | TauC | TauJ |
|---|---|---|---|---|
| *ADA* | 1.06 ± 0.09 | 1.04 ± 0.06 | 3.47 ± 0.42* | 1.35 ± 0.24# |
| *ADK* | 1.02 ± 0.03 | 1.06 ± 0.02 | 1.17 ± 0.06 | 0.85 ± 0.09# |
| *CD39* | 1.01 ± 0.03 | 1.23 ± 0.02* | 1.45 ± 0.06* | 0.94 ± 0.05# |
| *CD73* | 1.02 ± 0.04 | 1.23 ± 0.07* | 1.33 ± 0.05* | 1.13 ± 0.05# |
| *ENT1* | 1.06 ± 0.07 | 0.93 ± 0.05 | 1.11 ± 0.08 | 1.04 ± 0.10 |
| *ENT2* | 1.02 ± 0.04 | 1.07 ± 0.04 | 1.13 ± 0.03 | 0.96 ± 0.04# |
| *ENT3* | 1.02 ± 0.04 | 1.13 ± 0.06 | 1.24 ± 0.07* | 0.91 ± 0.06# |
| *ENT4* | 1.01 ± 0.04 | 1.15 ± 0.05 | 1.15 ± 0.06 | 0.85 ± 0.06# |

Mice were treated as indicated (control WT mice, WTC; J4-treated WT mice, WTJ; control Tau22 mice, TauC; and J4-treated Tau22 mice, TauJ; n= 6-9 in each group) from the age of 3-11 months. The hippocampus was harvested carefully and subjected to RT-qPCR. GAPDH was used as a reference gene for normalization. The data are expressed as the mean ± SEM. *$p<0.05$ versus the WT vehicle group; #$p<0.05$ versus the Tau22 vehicle group.

**Supplementary Table 9.** **The upregulated DAAs genes in the hippocampus of Tau22 mice**

| Type | Gene name | TauC/WTC | *p*_value | TauJ/TauC | *p*_value |
|---|---|---|---|---|---|
| **DAA/GFAP low** (38/239 upregulated genes) | *Abca1* | **0.45*** | 0.00 | -0.02 | 0.86 |
| | *Aspg* | **0.81*** | 0.01 | -0.25 | 0.40 |
| | *B2m* | **0.33*** | 0.00 | -0.09 | 0.27 |
| | *C1qa* | **0.45*** | 0.00 | **-0.21#** | 0.01 |
| | *C4b* | **1.16*** | 0.00 | 0.00 | 0.99 |
| | *Cd151* | **0.40*** | 0.00 | **-0.54#** | 0.00 |
| | *Cd9* | **0.58*** | 0.00 | -0.12 | 0.15 |
| | *Ctsd* | **0.41*** | 0.00 | -0.13 | 0.10 |
| | *Ezr* | **0.44*** | 0.00 | **-0.37#** | 0.00 |
| | *Fxyd1* | **0.59*** | 0.00 | **-0.54#** | 0.00 |
| | *Gadd45g* | **0.34*** | 0.00 | **-0.40#** | 0.00 |
| | *Gfap* | **0.73*** | 0.00 | -0.14 | 0.22 |
| | *Ggta1* | **0.68*** | 0.00 | -0.13 | 0.40 |
| | *H2-D1* | **0.56*** | 0.00 | **-0.26#** | 0.01 |
| | *H2-K1* | **0.53*** | 0.00 | **-0.34#** | 0.00 |
| | *Id4* | **0.42*** | 0.00 | **-0.43#** | 0.00 |
| | *Lamp2* | **0.37*** | 0.00 | **-0.19#** | 0.03 |
| | *Lgals3bp* | **0.92*** | 0.00 | **-0.39#** | 0.00 |
| | *Mgst1* | **0.49*** | 0.00 | **-0.40#** | 0.00 |
| | *Mt1* | **0.57*** | 0.00 | **-0.67#** | 0.00 |
| | *Mt2* | **0.81*** | 0.00 | **-0.72#** | 0.00 |
| | *Nfe2l2* | **0.57*** | 0.00 | **-0.34#** | 0.00 |
| | *Nfkbia* | **0.96*** | 0.00 | **-0.85#** | 0.00 |
| | *Osmr* | **0.56*** | 0.00 | 0.00 | 0.98 |
| | *Pard3b* | **0.48*** | 0.00 | -0.13 | 0.35 |
| | *Pdgfd* | **0.48*** | 0.00 | **-0.31#** | 0.03 |
| | *Pdzd2* | **0.40*** | 0.02 | **-0.39#** | 0.03 |
| | *Plce1* | **0.72*** | 0.00 | -0.15 | 0.31 |
| | *Sdc4* | **0.66*** | 0.00 | **-0.33#** | 0.00 |
| | *Serpina3n* | **0.52*** | 0.00 | 0.03 | 0.72 |
| | *Serpinf1* | **0.51*** | 0.00 | **-0.44#** | 0.01 |
| | *Serping1* | **0.46*** | 0.00 | **-0.49#** | 0.00 |
| | *Slc14a1* | **0.36*** | 0.00 | 0.10 | 0.28 |
| | *Stat3* | **0.37*** | 0.00 | -0.06 | 0.58 |
| | *Sulf1* | **1.97*** | 0.00 | **-1.66#** | 0.00 |
| | *Thbs4* | **0.40*** | 0.00 | -0.16 | 0.12 |
| | *Usp53* | **0.49*** | 0.00 | **-0.31#** | 0.00 |
| | *Vim* | **0.37*** | 0.00 | **-0.22#** | 0.03 |

The data are expressed as the log2 ratio. *$p<0.05$ versus the WT vehicle group; #$p<0.05$ versus the Tau22 vehicle group.

**Supplementary Table 9. The upregulated DAAs genes in the hippocampus of Tau22 mice (Continued)**

| Type | Gene name | TauC/WTC | *p*_value | TauJ/TauC | *p*_value |
|---|---|---|---|---|---|
| **GFAP high/GFAP low** (31/311 upregulated genes) | *Acsl3* | **0.36*** | 0.01 | -0.04 | 0.79 |
| | *Ass1* | **0.35*** | 0.00 | **-0.33#** | 0.00 |
| | *B2m* | **0.33*** | 0.00 | -0.09 | 0.27 |
| | *C4b* | **1.16*** | 0.00 | 0.00 | 0.99 |
| | *Cd9* | **0.58*** | 0.00 | -0.12 | 0.15 |
| | *Ctsd* | **0.41*** | 0.00 | -0.13 | 0.10 |
| | *Enpp2* | **2.25*** | 0.00 | **-2.03#** | 0.00 |
| | *Eva1a* | **0.37*** | 0.00 | **-0.50#** | 0.00 |
| | *Ezr* | **0.44*** | 0.00 | **-0.37#** | 0.00 |
| | *Fxyd1* | **0.59*** | 0.00 | **-0.54#** | 0.00 |
| | *Gadd45g* | **0.34*** | 0.00 | **-0.40#** | 0.00 |
| | *Gfap* | **0.73*** | 0.00 | -0.14 | 0.22 |
| | *Ggta1* | **0.68*** | 0.00 | -0.13 | 0.40 |
| | *H2-D1* | **0.56*** | 0.00 | **-0.26#** | 0.01 |
| | *H2-K1* | **0.53*** | 0.00 | **-0.34#** | 0.00 |
| | *Hhatl* | **0.33*** | 0.03 | -0.23 | 0.14 |
| | *Id4* | **0.42*** | 0.00 | **-0.43#** | 0.00 |
| | *Lamp2* | **0.37*** | 0.00 | **-0.19#** | 0.03 |
| | *Mdk* | **0.63*** | 0.00 | **-0.69#** | 0.00 |
| | *Mgst1* | **0.49*** | 0.00 | **-0.40#** | 0.00 |
| | *Msx1* | **1.35*** | 0.00 | **-1.38#** | 0.00 |
| | *Mt1* | **0.57*** | 0.00 | **-0.67#** | 0.00 |
| | *Mt2* | **0.81*** | 0.00 | **-0.72#** | 0.00 |
| | *Myoc* | **0.34*** | 0.00 | **-0.24#** | 0.01 |
| | *Nfe2l2* | **0.57*** | 0.00 | **-0.34#** | 0.00 |
| | *Nfkbia* | **0.96*** | 0.00 | **-0.85#** | 0.00 |
| | *Pdgfd* | **0.48*** | 0.00 | **-0.31#** | 0.03 |
| | *Sdc4* | **0.66*** | 0.00 | **-0.33#** | 0.00 |
| | *Serpinf1* | **0.51*** | 0.00 | **-0.44#** | 0.01 |
| | *Slc14a1* | **0.36*** | 0.00 | 0.10 | 0.28 |
| | *Vim* | **0.37*** | 0.00 | **-0.22#** | 0.03 |

The data are expressed as the log2 ratio. **p*<0.05 versus the WT vehicle group; #*p*<0.05 versus the Tau22 vehicle group.

### Supplemental Figures and Legends

#### Supplementary Figure 1

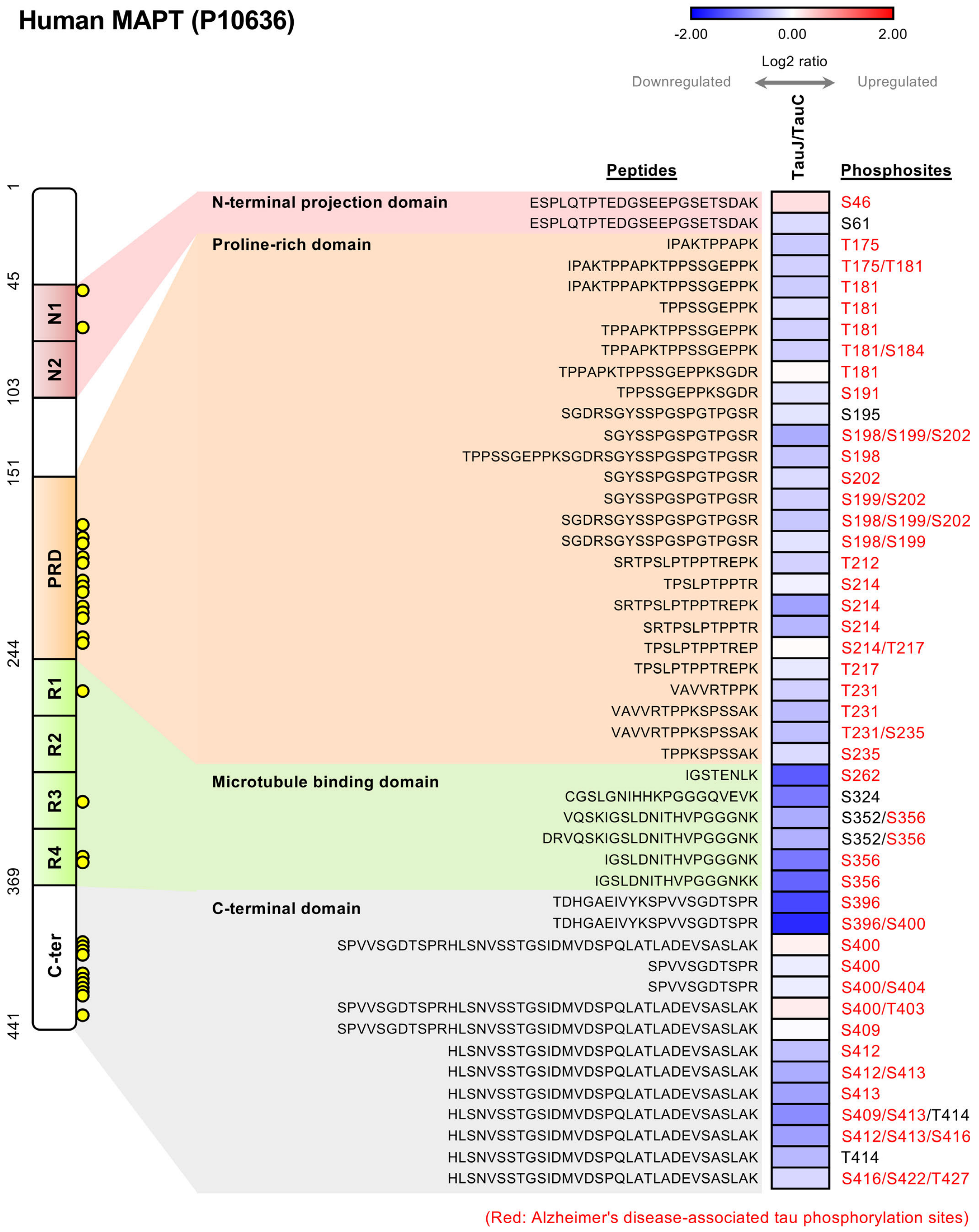


**Supplementary Fig. 1 Chronic J4 treatment reduces hyperphosphorylation at multiple sites in human Tau in the hippocampi of Tau22 mice**

Mice were treated as indicated (control Tau22 mice, TauC; J4-treated Tau22 mice, TauJ; n= 3 in each group) from the age of 3-12 months. Pooled total hippocampal lysates (200 μg) from 3 animals of the age of 12 months were harvested and subjected to phospho-proteomic analysis. The heatmap shows the relative log2 expression ratio of phosphorylated human tau (MAPT, P10636) in the TauJ group vs. the TauC group. The relative expression level (log2 ratio) of human phosphorylated tau is shown on a scale from red (upregulated) to blue (downregulated). Alzheimer's disease-associated tau phosphorylation sites are shown in red.

#### Supplementary Figure 2

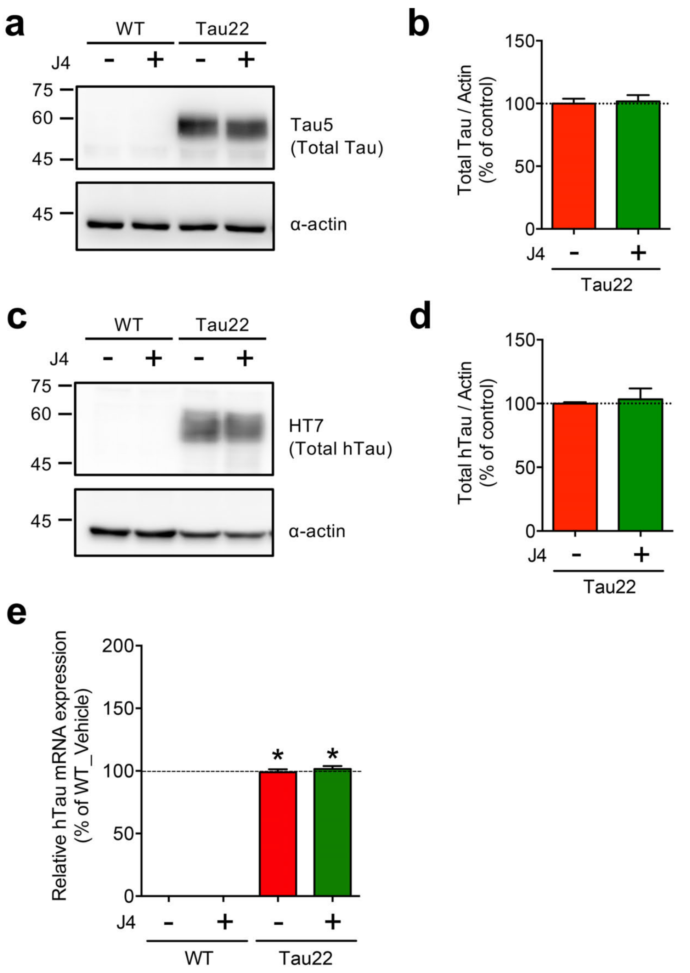


**Supplementary Fig. 2 Chronic J4 treatment has no effect on the expression of the human Tau protein and transgene in the hippocampi of Tau22 mice**

Mice were treated as indicated (control WT mice, WTC; J4-treated WT mice, WTJ; control Tau22 mice, TauC; and J4-treated Tau22 mice, TauJ n = 3 - 4 in each group) from the age of 3-11 months. The expression levels of **(a)** tau protein (Tau5) and **(c)** human tau protein (HT7) were detected using the indicated antibodies. α-Actin was used as a loading control. The protein expression level and the phosphorylation level were quantified and are shown in **(b)** and **(d)**. **(e)** The gene expression level of human Tau in mice from the different treatment groups (WTC, black; WTJ, blue; TauC, red; TauJ, green; n = 3-5) from the age of 3-11 months was examined by RT-qPCR. The data are expressed as the mean ± S.E.M. *$p < 0.05$ versus the WTC group; one-way ANOVA.

**Supplementary Figure 3**

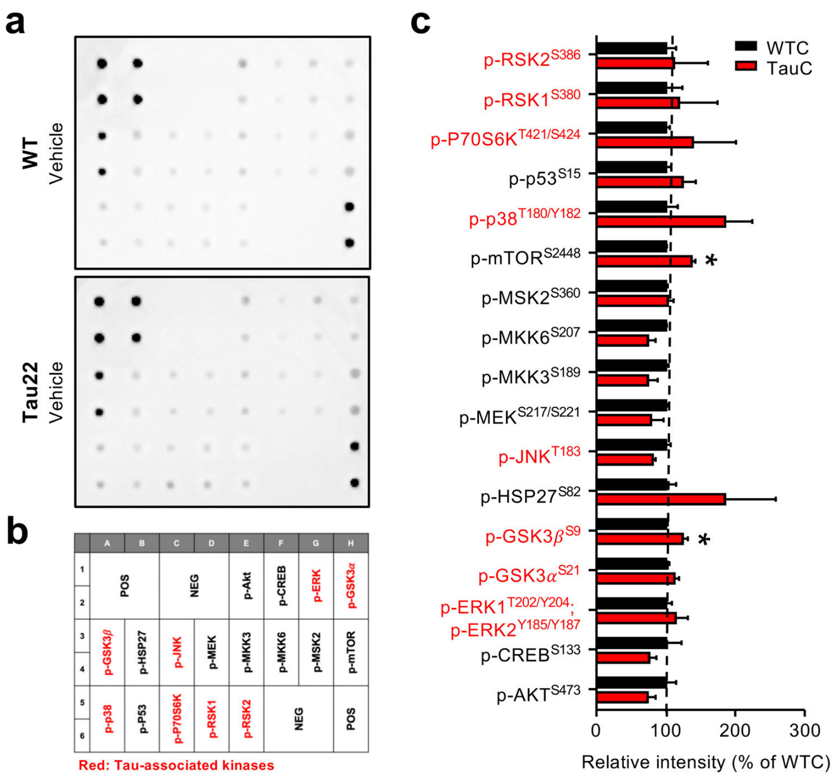


**Supplementary Fig. 3 MAPK phosphorylation array analysis of Tau22 mice.**

Mice were treated as indicated (control WT mice, WTC; and Control Tau22 mice, TauC; n = 6 in each group) from the age of 3-11 months. Pooled total hippocampal lysates (1 mg) were harvested and subjected to MAPK phosphorylation array. **(a)** Phosphorylation level of MAP kinases (including Tau-associated kinases) was assessed following the manufacturer's protocol. **(b)** The map of MAPK phosphorylation array. Words in red are Tau-associated kinases. Data from each array was normalized to the averaged positive control signals (POS) followed by being normalized to the WTC group. The relative intensity of indicated kinase is shown in **(c)**. The data are expressed as the mean ± S.E.M. *$p<0.05$ versus the WTC group, two-tailed Student's *t*-test.

**Supplementary Figure 4**

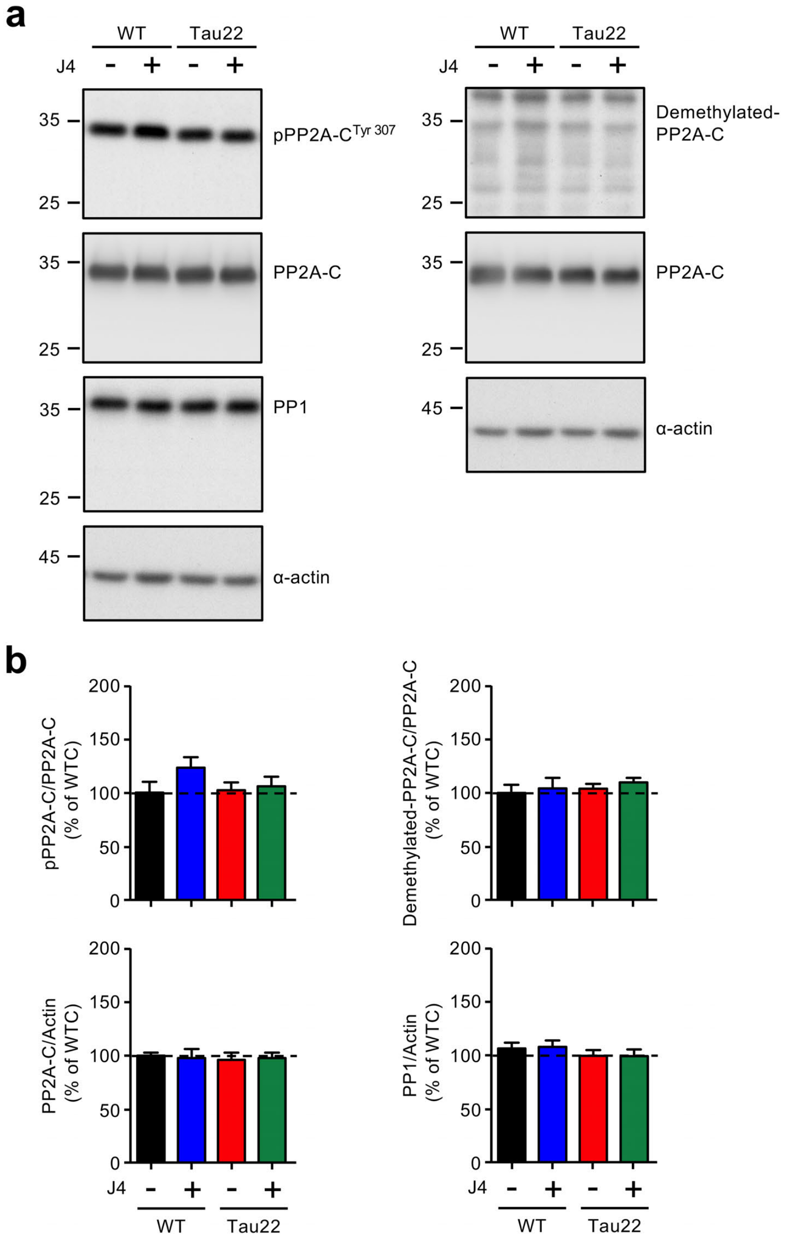


**Supplementary Fig. 4 Activities and expressions of PP1 and PP2A in Tau22 mice.**
Mice were treated as indicated (control WT mice, WTC, black; J4-treated WT mice, WTJ, bue; control Tau22 mice, TauC, red; and J4-treated Tau22 mice, TauJ, green; n = 5-7 in each group) from the age of 3-11 months. The total hippocampal lysates were harvested carefully, and subjected to Western blot analysis (30 μg per lane) for the amounts of phospho-PP2A-C (pTyr307), demethylated-PP2A-C, PP2A, and PP1 as indicated. The level of the indicated signal was normalized to that of α-Actin (a loading control) and shown in **(b)**. The data are expressed as the mean ± S.E.M.

Supplementary Figure 5

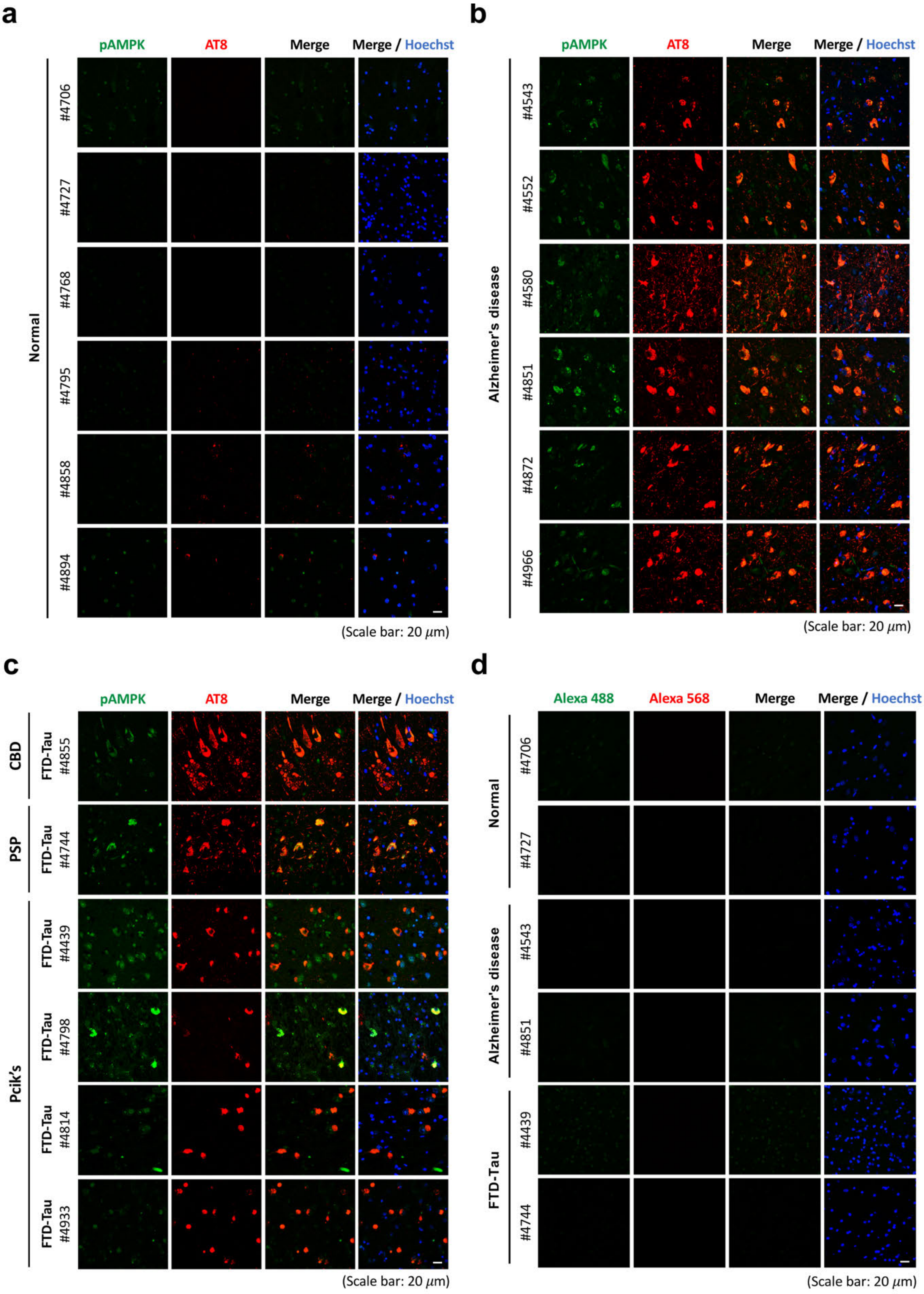


**Supplementary Fig. 5 AMPK activation can be observed in the posterior hippocampus of Alzheimer's disease and FTD-Tau patients.**

Posterior hippocampal sections (6 μm) from six normal subjects **(a)**, six Alzheimer's disease patients **(b)**, and six FTD-Tau (CBD, PSP, and Pick's) patients **(c)** were subjected to IHC staining. The level of phospho-AMPK and hyperphosphorylation tau was evaluated by staining with the indicated antibodies (pAMPK$^{Thr172}$, green; AT8 for pTau$^{Ser202/Thr205}$, red). The negative control is shown in **(d)**. Scale bar, 20 μm.

#### Supplementary Figure 6

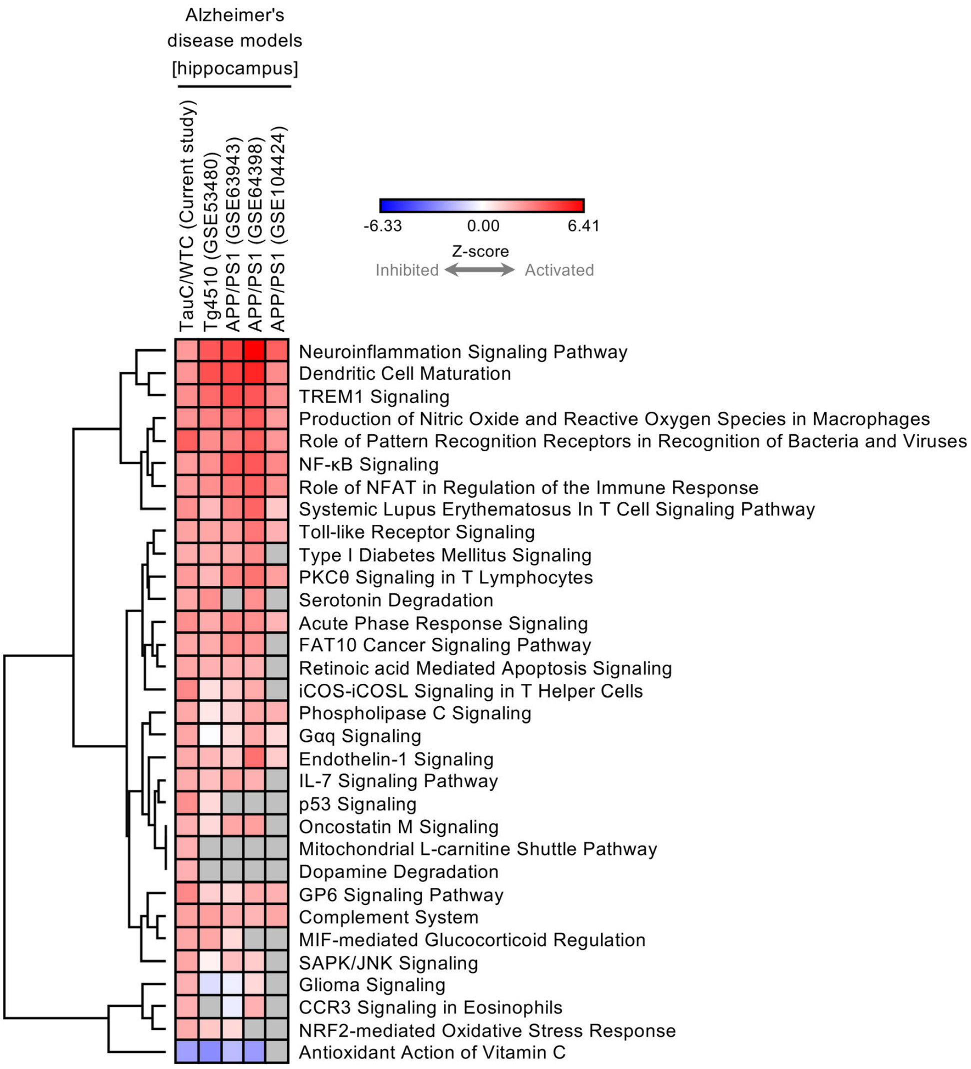


**Supplementary Fig. 6 Comparable canonical pathways are observed among Tau22 mice and other Alzheimer's disease mouse models**

The heatmap shows the canonical pathways enriched in the hippocampus of Tau22 mice and other Alzheimer's disease mouse models (Tg4510, GSE53480; APP/PS1, GSE63943/GSE64398/ GSE104424). The 1441 DE genes of the TauC/WTC group were subjected to IPA-Analysis Match analysis. Z-score is used to predict activation (Z-score ≥ 2.0, red) or inhibition (Z-score ≤− 2.0, blue) of regulation signaling.

**Supplementary Figure 7**

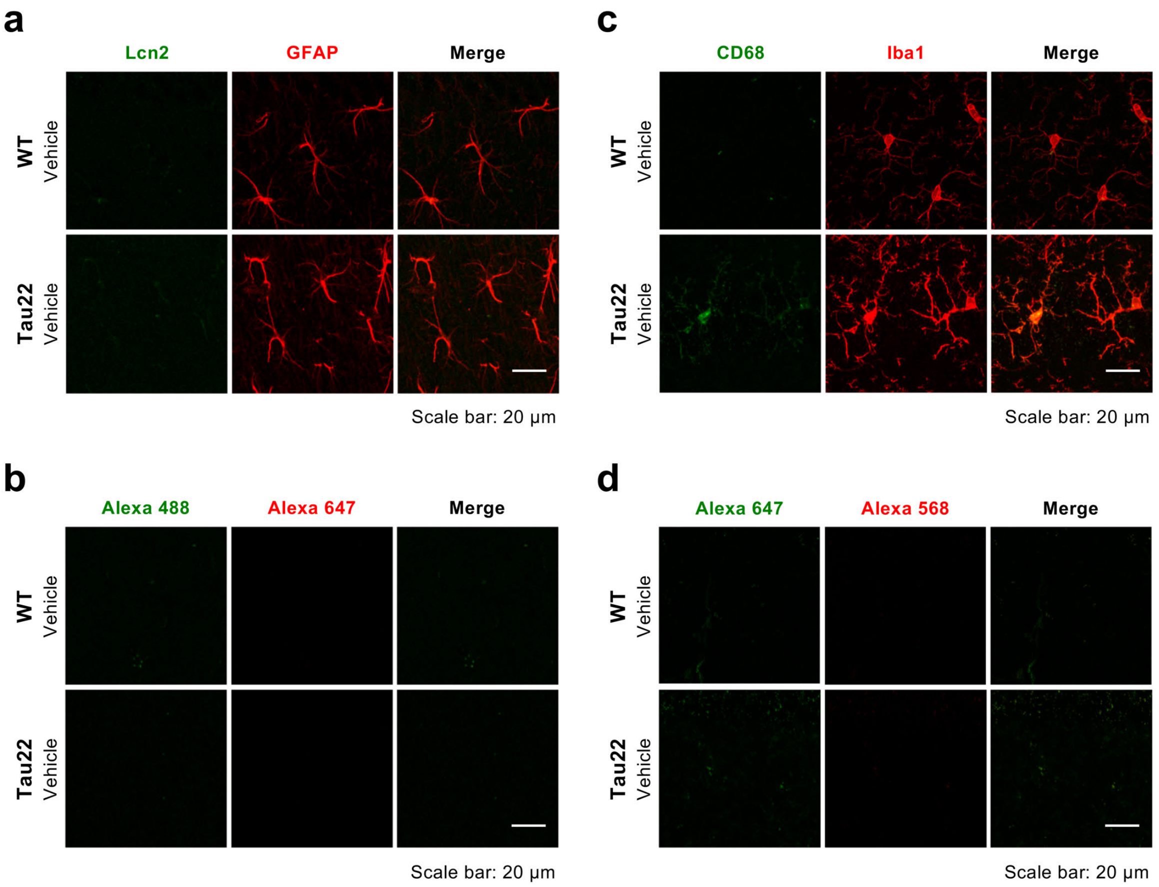


**Supplementary Fig. 7 Microgliosis, but not astrogliosis can be observed at 4 months of age in the hippocampus of THY-Tau22 mice.**

Hippocampal sections (20 μm) were prepared from Tau22 mice (n=3) and their littermate control (n=3) at the age of 4 months and subjected to IHC staining. The level and the activity of the **(a)** astrocyte and **(c)** microglia in the hippocampus were evaluated by staining with the indicated antibodies (GFAP for astrocyte, red; Lcn2 for reactive astrocyte, green; Iba1 for microglia, red; CD68 for reactive microglia, green), and the negative control are shown in **(b and d)**. Scale bar, 20 μm.

**Supplementary Figure 8**

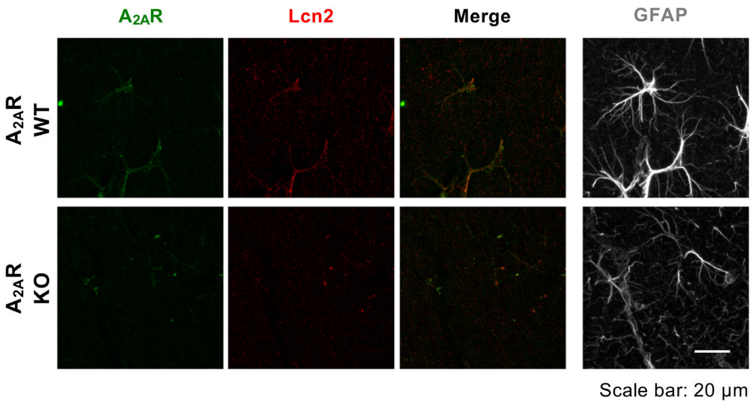


**Supplementary Fig. 8 $A_{2A}R$ and Lcn2 expressions are absent in the hippocampus of $A_{2A}R$ KO mice.** Hippocampal sections (20 μm) of $A_{2A}R$ WT and KO mice were used as a positive and negative control for the $A_{2A}R$ antibody. The level of $A_{2A}R$-positive astrocyte in the hippocampus was evaluated by staining with the indicated antibodies (anti-$A_{2A}R$ for $A_{2A}$ receptor, green; Lcn2 for reactive astrocyte, red; GFAP for astrocyte, grey). Scale bar, 20 μm.

Supplementary Figure 9

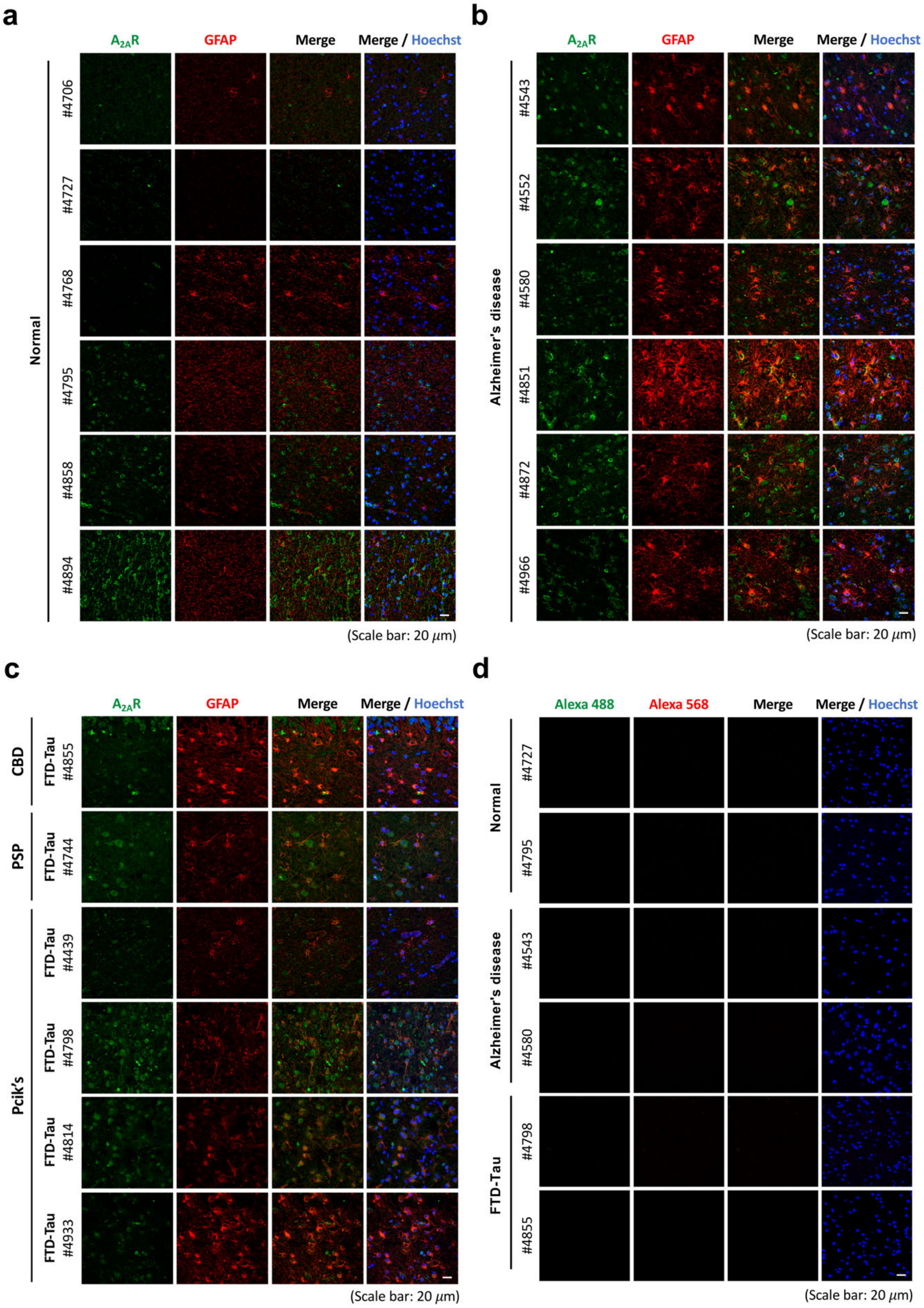


**Supplementary Fig. 9 Detection of astrocytic $A_{2A}R$ in the posterior hippocampus of patients with Alzheimer's disease or FTD-Tau, but not in normal subjects.**

Posterior hippocampal sections (6 μm) from six normal subjects **(a)**, six Alzheimer's disease patients **(b)**, and six FTD-Tau (CBD, PSP, and Pick's) patients **(c)** were subjected to IHC staining. he levels of $A_{2A}R$ and GFAP were evaluated by staining with the indicated antibodies (human $A_{2A}R$, green; GFAP, red). The negative control is shown in **(d)**. Scale bar, 20 μm.

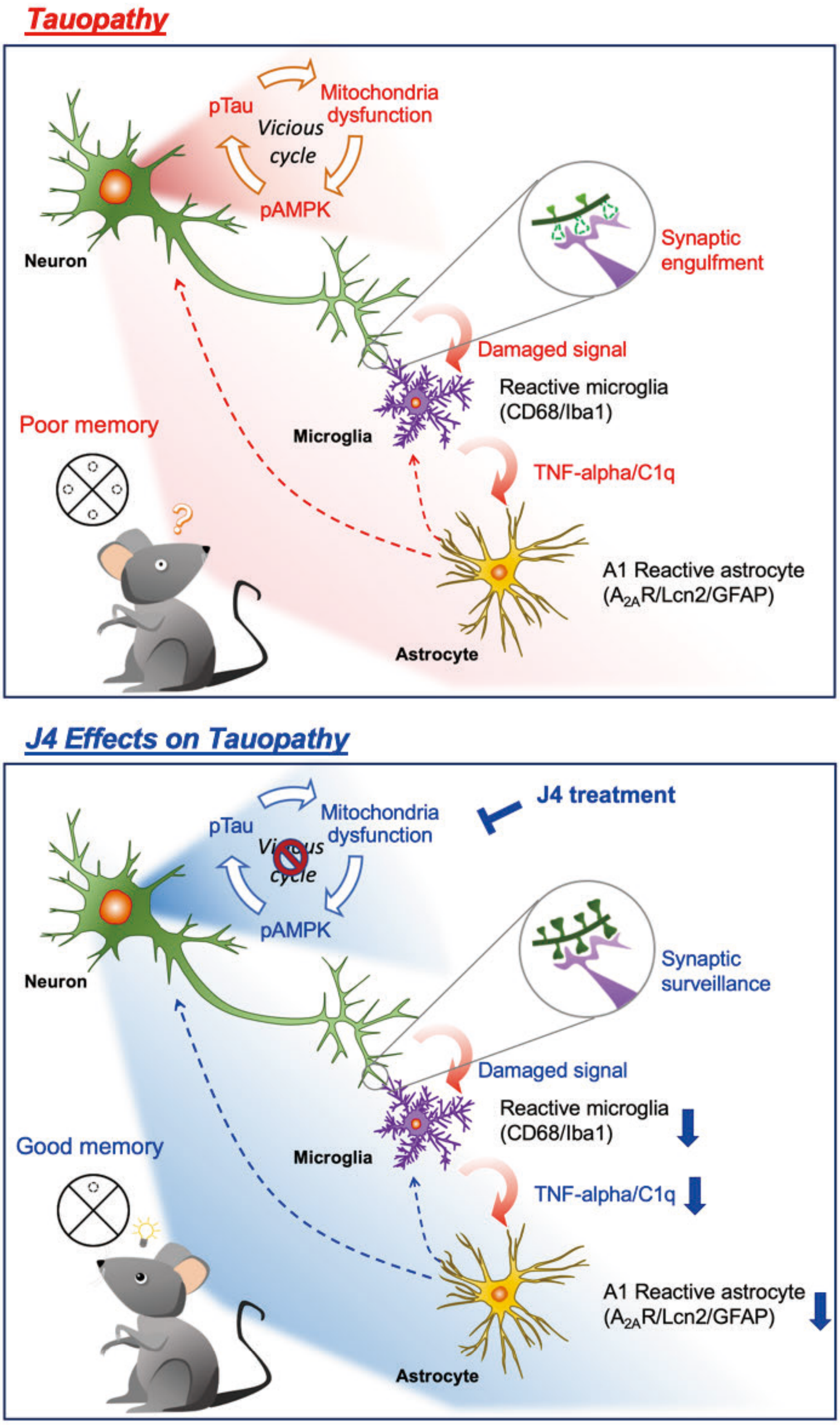


**Supplementary Fig. 10 Graphic abstract**

#### Supplemental Experimental Procedures

##### Morris water maze

Spatial memory and cognitive flexibility were evaluated using the Morris water maze test as described with slight modifications (Chang *et al.*, 2016). A circular swimming pool (154 cm in diameter, 51 cm in height) was filled with milky water (30 cm in depth, kept at 20 °C), and divided into four quadrants (T: target; L: target left; R: target right; O: opposite) with distinct visual cues on the tank wall of each quadrant. The hidden platform (13 cm in diameter, 0.5 cm below the surface of the milky water) was placed in the center of the target quadrant (T). Each mouse underwent four daily training trial (120 s/trial, 30 min interval), in which they were released from randomly selected nontarget quadrants (NTs). For the spatial memory test in the acquisition-learning phase, learning trials were performed with a hidden platform for five consecutive days (Day1-Day5). For the spatial reversal memory test in the reversal-learning phase, learning trials were performed with a hidden platform relocated to the opposite quadrant for an additional four consecutive days (Day 9-Day12). To evaluate reference memory, the probe trial and reversal probe trial were performed on Day 8 (72 hrs after the acquisition-learning phase) and Day 15 (72 hrs after the reversal-learning phase), respectively. For the probe test, the hidden platform was first removed. The mouse was then released in the opposite quadrant (O), and their swimming path was recorded for 120 s. The swimming path and other parameters (e.g. escape latency and swimming speed) of each mouse in different quadrants were monitored and analyzed using the TrackMot video tracking system (Diagnostic & Research Instruments Co., Ltd., Taoyuan, Taiwan). Mice that exhibited nonsearching behaviors (floating, a swimming speed below 10 cm/s and circling) were excluded from the analysis. Statistical differences were analyzed by two-way ANOVA.

##### Electrophysiological study

Mice aged 10-11 months were used for electrophysiology approaches. All electrophysiology studies were performed at the electrophysiology core facility (Neuroscience Program of Academia Sinica, Taipei, Taiwan). After rapid decapitation, the hippocampus was quickly dissected out and immersed in ice-cold artificial cerebrospinal fluid (ACSF; 119 mM NaCl, 2.5 mM KCl, 2.5 mM $CaCl_2$, 1.3 mM $MgSO_4$, 1 mM $NaH_2PO_4$, 26.2 mM $NaHCO_3$, and 11 mM glucose) oxygenated with 95% $O_2$ and 5% $CO_2$. Transverse hippocampal slices of 450 μm thickness were prepared with a DSK Microslicer (DTK-1000, Osaka, Japan) filled with oxygenated ice-cold ACSF. For recovery, the slices were then incubated in an interface-type holding chamber filled with oxygenated ACSF at RT for at least 3 hrs. Before recording, the recording electrodes were prepared with a glass micropipette puller (PC-10, Narishige, Tokyo, Japan), and the slices were transferred to an immersion-type recording chamber equipped with a perfusion system (flow rate: 2-3 ml/min) and temperature controller (kept at 32 °C). To record field excitatory postsynaptic potentials (fEPSPs) recording, the bipolar stainless-steel stimulating electrodes (Frederick Haer Company, Bowdoinham, ME; 10 ΩM impedance) and a glass pipette filled with 3 M NaCl were placed in the stratum radiatum of the hippocampal CA1 region. Basal synaptic transmission at the Schaffer collateral-CA1 synapses was first evaluated by measuring input-output curves using 12 stimuli (constant current pulses from 10 μA to 120 μA in increments of 10 μA, duration of 40 μs). To measure the paired-pulse facilitation (PPF) response, two pulses were applied in rapid succession (interpulse intervals of 50, 100, 150, 200, 300, 400 and 500 ms). Baseline responses were recorded by applying single stimuli (40 µsec pulse-width) at 30s intervals, and 2 responses were averaged to obtain a data point. Long-term depression (LTD) was induced using 3 trains of low-frequency stimulation (LFS, 1200 pulses at 2 Hz) with a 10-min interval/train as described elsewhere (Van der Jeugd *et al.*, 2011). The initial fEPSP slope was calculated by using Signal software (V4.08, Cambridge Electronic Design, Cambridge, UK).

##### Immunoblotting

After the animals were rapidly decapitated, the hippocampus was dissected out, homogenized in an ice-cold detergent-free lysis buffer (10 mM Tris-HCl, pH 7.4, and 0.32 M sucrose) containing inhibitors of protease (50X complete EDTA free Tab, Roche Diagnostics, IN, USA) and phosphatase inhibitors (10x PhosSTOP, Roche Diagnostics) using sterilized tissue grinder (FocusBio, Taiwan) and then passed through an insulin syringe 15 times. The tissue lysates were then centrifuged at 800 x g for 10 min at 4 °C to remove the debris. The supernatants were collected and stored at -80 °C until use. The protein concentration was measured using the Bio-Rad Protein Assay Dye Reagent Concentrate (Bio-Rad, Hercules, CA, USA). For Western blot analyses, total proteins (20 µg) were first separated via SDS-PAGE (10%-12%), transferred onto PVDF membranes (0.45 µm pore size, Millipore, MA, USA), hybridized with the indicated primary antibody (Table S4) overnight at 4 °C, and then incubated with the corresponding horseradish peroxidase (HRP)-conjugated secondary antibody for 1 hr at RT. After extensive washes, the immunosignals were detected using enhanced chemiluminescence (ECL) reagent (PerkinElmer, MA, USA).

##### Phospho-proteomic analysis

Hippocampal lysates (200 µg) collected from three animals from each condition were subjected to phospho-proteomic analysis at the Proteomics Core Facility (PCF) (Institute of Biomedical Sciences, Academia Sinica, Taipei, Taiwan). For in-solution digestion, 8 M urea was added to the lysates to prepare a mixture of 10 mg protein/ml in 6 M urea. Protein reduction was performed by the addition of dithiothreitol (DTT, final concentration 5 mM) and incubation at 56 °C for 25 min. Protein alkylation was performed by the addition of iodoacetamide (IAA, final concentration 15 mM) and incubation at RT for 45 mins to block the reversion of sulfhydryl (-SH) groups to disulfide bonds. Trypsin/Lys-C Mix (Promega, WI, USA) was then added to protein (protein: protease = 25:1 w/w) and the mixture was incubated for 4 hrs at 37 °C. The urea concentration was adjusted to 1 M or less by diluting the reaction mixture with triethylammonium bicarbonate (TEAB, 50 mM) followed by an incubation at 37 °C for 17 hrs. The digested samples were dried with a SpeedVac and desalted with C18 Oasis® PRiME HLB cartridges (Waters, MA, USA). For iTRAQ labeling, the digested peptides were labeled with four isobaric iTRAQ Reagents (114, 115, 116, and 117) using iTRAQ® Reagents-4plex Applications Kit (AB Sciex, MA, USA) following manufacturer's instructions. For phosphopeptide enrichment, iTRAQ labeled peptides were mixed with loading buffer (80% acetonitrile, ACN, 5% trifluoroacetic acid, TFA, and 1 M glycolic acid) and adjusted to pH 2. The sample solution was then mixed with $TiO_2$ beads (GL Sciences, Japan) and incubated with vortexing at RT for 15 mins. The beads were collected and washed twice with 100 µl washing buffer (80% ACN and 5% TFA). The phosphopeptides were sequentially eluted with 50 µl 0.5% $NH_4OH$, 50 µl 5% $NH_4OH$, and 50 µl 80% ACN with 0.1% formic acid, and dried with a SpeedVac. The phosphopeptides were then subjected to LC/MS/MS and analyzed by Proteome Discoverer ver.2.2 (Thermo Fisher Scientific, Waltham, MA, USA).

#### Phospho-kinase array

Mouse hippocampal lysates collected from three animals from each condition were analyzed with the Human/Mouse MAPK Phosphorylation Array (C-Series, Cat# AAH-MAPK-1-4, RayBiotech Inc., USA) according to the manufacturer's protocol. The samples (1 mg/ml in the blocking buffer) were loaded onto a membrane that preincubated with the blocking buffer for 30 min at RT and incubated overnight at 4 °C. The membrane was then washed with wash buffer I and wash buffer II for 3 times each and then incubated with a detection antibody cocktail overnight at 4 °C. After extensive wishes, the membrane was incubated with HRP-conjugated anti-rabbit IgG overnight at 4 °C and washed extensively. The immunosignals were monitored using chemiluminescent detection buffer. The intensity of each spot on the membrane was quantified with ImageJ software (NIH, Bethesda, MD, USA) and normalized to the intensities of the positive control on the same membrane. The relative kinase activity was assessed by normalizing the signal of the phosphorylated substrate to that of the positive controls.

#### RNA extraction, cDNA synthesis, and quantitative PCR

For mouse brain tissue, RNA isolation and complementary DNA (cDNA) synthesis were performed according to the manufacturer's protocols. In brief, mouse hippocampal tissues were homogenized in GENEzol™ reagent (GZX100, Geneaid Biotech Ltd., New Taipei City, Taiwan) with sterilized tissue grinders (FocusBio, Taiwan), and then standard procedures for RNA preparation and cDNA synthesis were performed as described previously (Lee *et al.*, 2018). Quantitative PCR (qPCR) assays were carried out using the LightCycler® 480 System (Roche Life Science, Indiana, USA) and analyzed by the comparative CT (ΔΔCt) method with GAPDH as a reference gene. The sequences of the PCR primers are shown in Table S2.

For Human brain tissue, total mRNA was extracted and purified using the RNeasyLipid Tissue Mini Kit (Qiagen). One microgram of total mRNA was reverse-transcribed using the HighCapacity cDNA reverse transcription kit (Applied Biosystems). Quantitative real-time polymerase chain reaction (qPCR) analysis was performed on an Applied Biosystems™ StepOnePlus™ Real-Time PCR Systems using TaqMan™ Gene Expression Master Mix (Applied Biosystems™). The thermal cycler conditions were as follows: 95°C for 10min, then 40 cycles at 95°C for 15 seconds and 60°C for 1 minute. Amplifications were carried out in duplicate and the relative expression of target genes was determined by the ΔΔCt method with □-Actin (ACTB) was used as a reference housekeeping gene for normalization. References of the probes used in this study are given in Table S3.

#### RNA sequencing (RNA-seq)

Total RNA samples (3 µg per sample) extracted from the hippocampus with RIN values greater than 8 were subjected to RNA-seq analysis. The RNA library construction and sequencing were carried out by Welgene Biotech (Taipei, Taiwan). Briefly, the SureSelect Strand-Specific RNA Library Preparation Kit (Agilent Technology, CA, USA) was used for library construction on the Illumina platform. After AMPure XP Bead-based (Beckman Coulter Genomics, MA, USA) size selection of the RNA library, the sequences were determined using the Illumina's sequencing-by-synthesis (SBS) technology to obtain 150-bp paired-end reads. Sequencing data (FASTQ files) were generated by Welgene's pipeline (Base call conversion, adaptor clipping, and sequence quality trimming) based on Illumina's base-calling program bcl2fastq v2.2.0 (Illumina, CA, USA) and Trimmomatic v0.36 (Bolger *et al.*, 2014). The RNA-seq reads were then aligned to the mouse reference genome (mm10) from the Ensembl database (Ensembl release 93) using HISAT2 (Kim *et al.*, 2015). Expression levels (fragments per kilobase per million, FPKM) were analyzed and estimated using cuffdiff (cufflinks v2.2.1) (Trapnell *et al.*, 2012) and Welgene in-house programs. To identify the differentially expressed (DE) genes in different groups, cutoff criteria (absolute $\log_2$ fold change $\geq 0.32$, $p<0.05$) were used. A volcano plot and heatmap of the DE genes were drawn by using Instant Clue (Nolte *et al.*, 2018) and Morpheus software (https://software.broadinstitute.org/ morpheus/), respectively. The Gene Ontology (GO) and the Kyoto Encyclopedia of Genes and Genomes (KEGG) pathways of DE genes were analyzed by using the Database for Annotation, Visualization and Integrated Discovery (DAVID 6.7) (Huang da *et al.*, 2009a, b). Ingenuity Pathway Analysis (IPA) software (Qiagen, CA, USA) was then used for the identification of signaling pathway(s) or upstream regulator(s) related to the DE genes.

##### Immunohistochemical staining

Coronal brain sections (20 μm) from the desired mice were prepared as previously described (Lee *et al.*, 2018). For immunofluorescence (IF) staining, brain slices were washed with 0.1 M PBS buffer, permeabilized with 0.2% Triton X-100 solution (in 0.1 M PBS buffer), and blocked with 3% normal goat serum (NGS), 3% normal donkey serum (NDS), or 3% bovine serum albumin (BSA) in 0.1 M PBS buffer for 2 hrs at RT. The brain sections were then washed with 0.1 M PBS buffer twice and incubated with the indicated primary antibodies (listed in Table S4) in primary antibody solution (1% NGS or BSA, 0.2% Triton X-100, and 0.1% sodium azide in 0.1 M PBS) for 48 hrs at 4 °C. After extensive washes, the brain sections were incubated with the corresponding secondary antibody (1:500) for 2 hrs at RT, and then the nuclei were stained with Hoechst 33258 (1:5000) for 10 min at RT. Free-floating brain sections were mounted on the silane-coated slides (Muto Pure Chemicals Co., Tokyo, Japan) with mounting media (Vector Laboratories, CA, USA) and stored at 4 °C before imaging. An LSM 780 confocal microscope (Carl Zeiss, Germany) was used to capture images. The images were analyzed with MetaMorph software (Universal Imaging, PA, USA).

Formalin-fixed, paraffin-embedded human brain slices were deparaffinized and rehydrated. To expose the antigenic sites, the brain slices were then immersed in 1X citrate buffer (C9999, Sigma-Aldrich, St. Louis, MO, USA) for 20 min at 97.5 °C and cooled to RT. The brain slices were subjected to immunofluorescence staining as described above. After secondary antibody (1:500) incubation, the brain sections were treated with 0.1% (w/v) Sudan Black B (199664, Sigma) in 70% ethanol for 15 min at RT to block autofluorescence signal. The brain sections were then stained with Hoechst 33258 (1:5000) for 10 min at RT. After extensive washes with PBS, the brain sections were mounted with the mounting media (Vector Laboratories, CA, USA) and stored at 4 °C before imaging.
